# Inherent coupling of perceptual judgments to actions in the mouse cortex

**DOI:** 10.1101/2025.10.30.685660

**Authors:** Michael Sokoletsky, Arina N. Nikitina, Kedar Waychal, Yonatan Katz, Ilan Lampl

**Author notes:** Equal contribution authors (Arina N. Nikitina, Kedar Waychal).

## Abstract

Neural activity during perceptual tasks is distributed across many cortical regions, but it remains unclear where perceptual judgments are made and whether they are encoded independently of the resulting actions. To address these questions, we designed a vibrotactile detection task in which mice flexibly switched between standard and reversed contingency blocks, requiring them to lick when a stimulus was present or absent, depending on the block. A cortex-wide optogenetic screen revealed that the premotor cortex is important for perceptual judgments rather than the ability to lick. However, coding of perceptual judgments in the premotor cortex and other cortical regions was coupled to actions rather than being independent of them. Finally, we identified a subset of premotor cortex cells whose activity encoded the current block identity. Based on these findings, we propose a model in which vibrotactile perceptual judgments are inherently, but flexibly, coupled to actions via cortical activity.

## Introduction

A key goal of neuroscience is to understand the cognitive processes that transform sensory stimuli into actions. Such transformations abound in our daily lives. For example, a fly that lands on our skin may cause us to shoo it off depending on whether we feel it. They can also be context-dependent—a yellow traffic light may lead us to accelerate or decelerate based on our proximity to the stop line and the presence of a car behind us. To study such transformations, a common approach is to train subjects on tasks that link stimuli to actions while recording or perturbing neural activity. This approach has been extensively applied to humans, non-human primates, and rodents, yielding key insights into sensorimotor transformations across sensory modalities and tasks, including tactile detection^1,2^, visual discrimination^3,4^, and olfactory matching^5^, among others.

Among these insights is that the neocortex plays a central role in the sensorimotor transformation; in fact, commonly proposed key regions—primary sensory^1,2,6^, higher-order sensory^7,8^, parietal^9–11^, retrosplenial^12–14^, prefrontal^15,16^, premotor^17–19^, and motor^20,21^—together span nearly the entire neocortex. Several subcortical regions, including the thalamus^22–24^, striatum^25–27^, and superior colliculus^28–30^, have also been proposed as key nodes. However, it remains unclear where and how the sensorimotor transformation takes place. In particular, some studies have suggested the sensorimotor transformation is mediated by neural activity that reflects perceptual judgments (i.e., the subject’s internal assessment of the stimulus) independently of the selected actions^1,16,31^. Others, by contrast, have posited that perceptual judgments are fundamentally coupled to actions, making such action-independent perceptual judgment coding superfluous^32–34^.

Searching for action-independent judgment coding has recently been complicated by the observation that at least in mice^35,36^ (and likely, to an extent, also in non-human primates^37^ and humans^38^), action-related activity appears throughout the brain, including in the primary sensory regions. As a result, activity in non-motor regions that is correlated with perceptual judgments may reflect feedback from the resulting actions rather than judgments per se^39^. This issue may be ameliorated by introducing a delay between stimulus and action, as well as by making animals perform symmetric actions depending on their judgment (e.g., licking left or right). However, recent studies have also observed widely distributed correlates of motor preparation^17,40^ and embodiment (i.e., uninstructed movements that predict the specific upcoming action)^41,42^, suggesting that both are incomplete solutions.

Here, we concentrate on a simple type of perceptual judgment, namely whether a vibrotactile whisker stimulus is present or absent (i.e., detection). To separate judgments from actions, we trained mice to switch between licking in the presence of the stimulus (standard block) and licking in its absence (reversed block), such that the same judgment is linked to different actions in each block. We found that whereas optogenetic inhibition of the somatosensory and premotor cortices impaired perceptual judgments to a much larger extent than it impaired actions, differences in neural activity across judgments in these areas were coupled to actions. At the same time, we found a subset of cells in the premotor cortex that encoded the current block in a transient manner. We created a neural network model in which this subset together with the stimulus input resulted in action-coupled perceptual judgments, reproducing the mouse’s behavior and key features of neural activity. Therefore, in mice performing vibrotactile detection, our results support a model in which the cortex maps stimulus inputs and task rules directly onto actions.

## Results

### Behaviorally decoupling perceptual judgments from actions

We trained mice (*n* = 25) to alternate between two blocks, called standard and reversed, in a vibrotactile detection task. In standard blocks, the mice had access to the left port while the right was retracted, and in reversed blocks, they had access to the right lick port while the left was retracted (Fig. 1a). Each trial began with a 100-ms, 5-kHz tone (Fig. 1b), which was identical across blocks. On half of the trials, a 100-ms, 20-Hz sinusoidal whisker stimulus was delivered to the left C2 whisker simultaneously with the tone. In standard blocks, mice needed to lick the left port when given the whisker stimulus and withhold licking when given the tone alone, while in reversed blocks, mice needed to lick the right port when given the tone alone and withhold licking when given the whisker stimulus. At the end of the tone, the mouse was presented with the non-retracted lick port and could lick it within a 500 ms response window. Hits within this window were rewarded with a drop of sucrose water, whereas false alarms were punished with a light airpuff. Rewards or punishments following correct or incorrect licks were provided immediately after the response window, regardless of lick timing.

**Figure 1:**
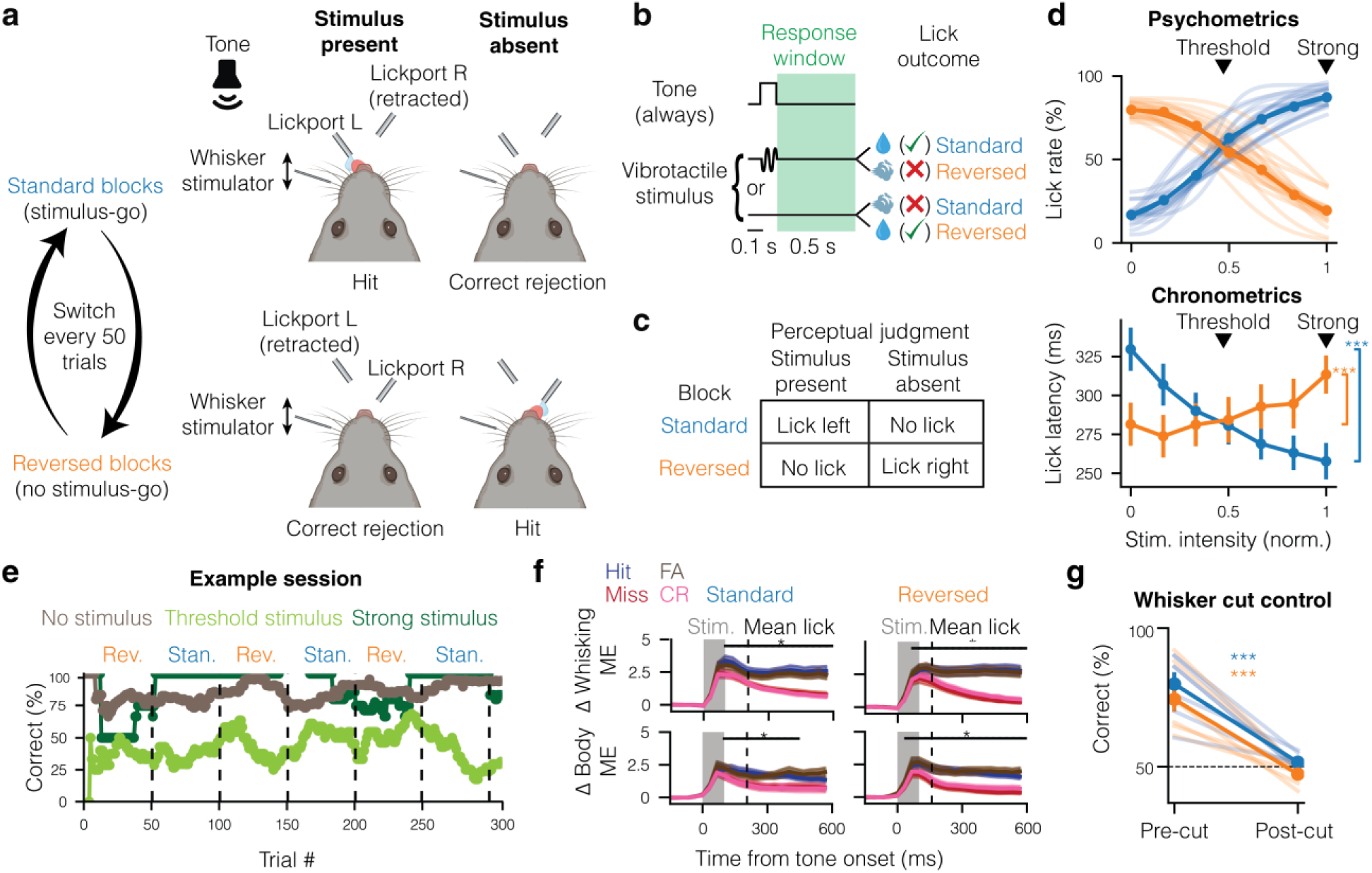
A contingency blocks task to separate perceptual judgments from actions. **(a)** Schematic of a mouse performing correct trials across stimulus present (left column) and stimulus absent (right column) trials on either standard blocks (top row) or reversed blocks (bottom row). The two blocks alternated once every 50 trials. **(b)** Schematic of trial structure in either block, starting with a 100-ms tone either by itself or together with a whisker stimulus, then a 500-ms response window, and then the outcome delivery, where licking the presented lick port leads to a sugar water reward or an airpuff punishment as outlined. **(c)** Table showing how perceptual judgments are directly coupled to actions in standard blocks, and inversely coupled in reversed blocks. Thus, by considering both blocks in conjunction, judgment and action become dissociable. **(d)** Psychometric curves (top) and chronometric curves (bottom) for each block, as a function of stimulus intensity (normalized to 1). **(e)** An example session of an expert mouse with correct responses to strong stimulus trials in olive green, threshold stimulus trials in light green, and no-stimulus trials in brown. Performance was smoothed with a 40-trial boxcar filter. Note that the green line reflects boxcar-smoothed performance over only 10% of trials in which the stimulus was strong, which yields 100%-correct periods during which the mouse responded correctly to each of the few strong stimulus trials, and a break near the middle, during which there were no strong stimulus trials. **(f)** Comparisons of changes in whisking (top panels) and body (bottom panels) motion energies (ME) relative to baseline in lick trials (hits and false alarms) and no-lick trials (misses and correct rejections) in both standard (left panels) and reversed (right panels) blocks. Significance was tested on the mean ME of lick vs. no-lick trials and corrected for multiple comparisons using the Dunn–Šidák correction. **(g)** Percent correct responses in each block before and after the C2 whisker was cut in the final session. Mean ± s.e.m. across *n* = 25 mice. * *P <* 0.05, ** *P <* 0.01, *** *P <* 0.001. Mouse illustration in (a) was partially created with BioRender.com.

Mice learned the task in two stages. In the first stage (Extended Data Fig. 1a,b), both lick ports were accessible, and mice were trained to lick left when given a strong (∼6° deflection) whisker stimulus together with the tone and lick right when given the tone alone. In the second stage, we began retracting one of the two lick ports every 50 trials, thereby implementing the block structure.

Block switches were signaled by the audible movement of the lick ports but were otherwise not cued deliberately. To assess switching, the first trial after each switch was a clear no-go trial (either no stimulus in the standard block or a strong stimulus in the reversed block). If the mice were still behaving as in the previous block, in which this would be a clear go trial, this would trigger an “air lick” towards the now out-of-reach lick port. In the initial block switches, air licks were frequent (∼75%) in the first post-switch trials and remained substantial (Extended Data Fig. 1c). After block training, however, air licks were rarely observed, appearing in only 15% of first post-switch trials and subsequently declining to nearly zero.

Given that mice were able to rapidly switch their behavior even before reinforcement, it is likely that they used a multimodal strategy to solve the task, combining the whisker stimulus input and tone with somatosensory and/or olfactory cues about the reachable lick port on each trial. Nonetheless, any successful strategy required the mouse to make perceptual judgments regarding whisker stimulus presence or absence (i.e., detection) during the tone period. The difference in relationship across blocks between the actions and inferred perceptual judgments then allowed us to decouple the two in trial-to-trial behavior (Fig. 1c, Extended Data Fig. 1d). Importantly, while behaviorally decoupling perceptual judgments from actions is a prerequisite for isolating action-independent judgment coding, it does not imply that such a code must necessarily exist (Supplementary Note 1, Supplementary Fig. 1).

After mice learned the task (*d’* > 1.5) with strong stimuli, we compared behavior across blocks with psychometric testing (Fig. 1d). As expected, mice licked more often as a function of stimulus intensity in standard blocks but less often in reversed blocks. Using model comparison (Methods; Extended Data Fig. 1e,f), we found that two logistic classifiers with different slopes and biases explained our curves most parsimoniously, with the magnitude of both slope and bias being slightly lower in reversed blocks (Extended Data Fig. 1g). Interestingly, chronometric (latency to lick) curves displayed the opposite trend to the psychometric curves across the blocks (Fig. 1d).

Subsequently, two classes of experiments were conducted—one in which we gave mice all stimulus intensities, and one in which we gave only threshold and strong stimuli. A representative example of the latter class is shown in Fig. 1e, where it can be observed that responses to no stimuli and strong stimuli were generally correct while responses to threshold stimuli fluctuated around 50%. We also recorded the mice’s movements using two cameras focused on their faces and bodies. Previous studies have shown that mice make uninstructed movements, such as whisking and limb movements, in a manner partially coupled to instructed movements like licking^35,42^. We replicated this result in both blocks (Fig. 1f), finding that licking in the reversed block was similarly associated with an increase in whisking and body motion as in the standard block, regardless of the trial type. This suggests that, across blocks, not only is the relationship between perceptual judgments and the instructed action (licking) reversed, but so is the relationship between perceptual judgments and uninstructed movements. Lastly, to confirm the mice’s reliance on the whisker stimulus, we cut the C2 whisker after the first two blocks on the final day of experiments. This reduced performance to chance level in both blocks (Fig. 1g).

### Dissociating causal effects on perceptual judgments and actions

As a first application of our approach, we wanted to assess which cortical regions are specifically important for perceptual judgments as opposed to actions. To do so, we used mice expressing ChR2 in inhibitory parvalbumin-positive (PV) cells, and transcranially inhibited cortical activity with a focused laser beam (Fig. 2a; *n* = 31,225 trials in total across 5 mice)^18,43,44^. In a control experiment in which we simultaneously recorded single-unit activity with a Neuropixels^45^ probe (*n* = 4 mice), we found the beam inhibited more than 80% of excitatory cell activity at the target, about 40% at 1.5 mm, and had a negligible effect at 3 mm (Extended Data Fig. 2a–d; 1.5 mm radius plotted in Fig. 2b). The laser was applied continuously from 200 ms before the tone or stimulus to the end of the response window (Fig. 2a). We did so rather than inhibiting during specific epochs to avoid assuming when the sensorimotor transformation took place. In a similar vein, to avoid assuming where in the cortex it took place, we conducted a global optogenetic screen at 42 locations across the dorsal cortical surface (Methods; Fig. 2b). These locations were aligned to the Allen Mouse CCF, with control laser delivery outside the cortex in 20% of trials. Stimuli were presented either at threshold or at strong intensity. We first examined the effect of laser inhibition on licking across all stimulus trials. While inhibiting the barrel cortex significantly reduced licking after the stimulus in standard blocks, as expected, it surprisingly had no effect in reversed blocks (Fig. 2c). In addition, we observed heterogeneous effects in other regions, such as how inhibiting the premotor cortex (M2; the most anterior region in Fig. 2c) in either hemisphere had an effect in the reversed blocks but inhibiting it only contralaterally had an effect in standard blocks.

**Figure 2:**
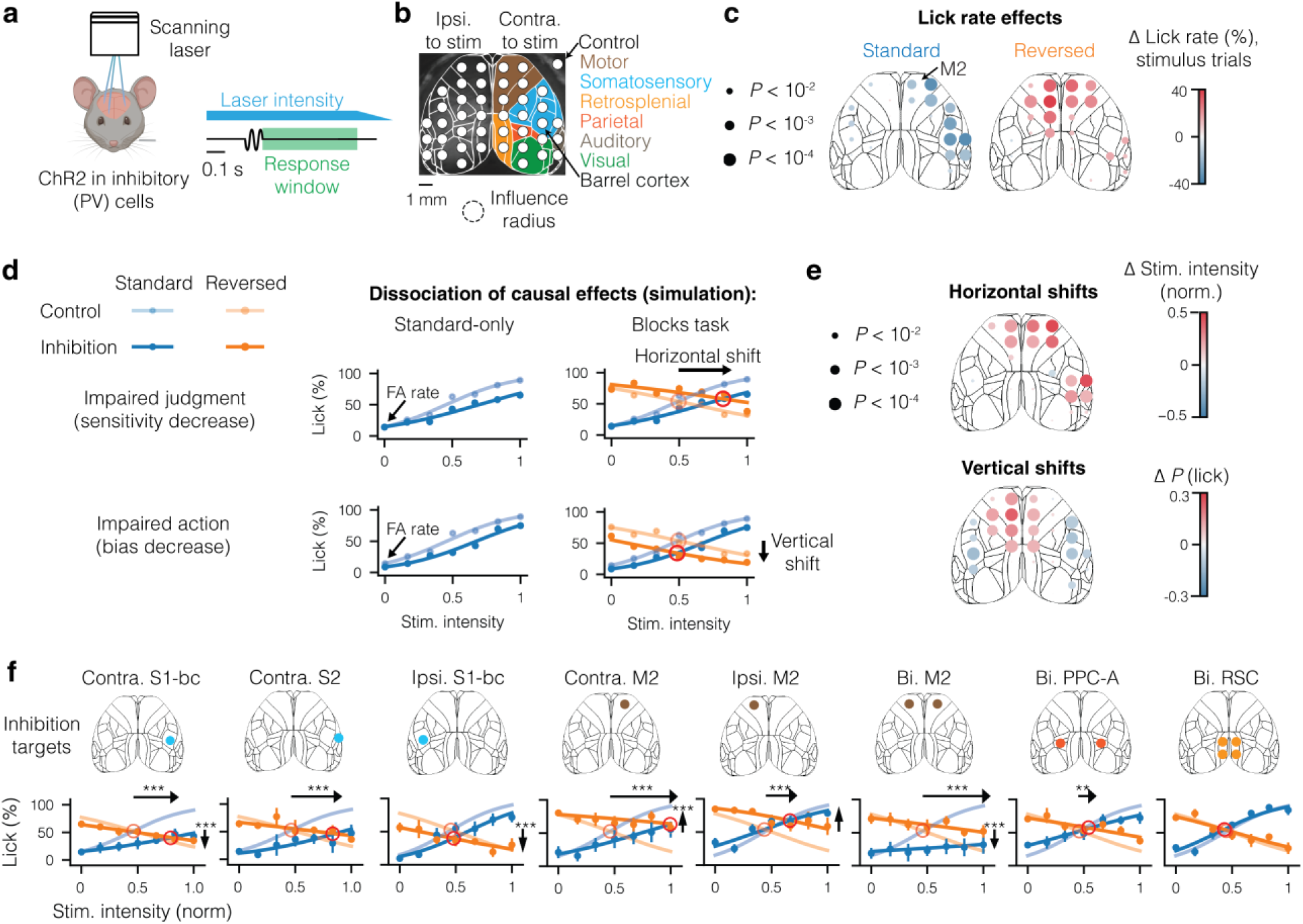
Contralateral somatosensory and premotor cortices are important for stimulus sensitivity. **(a)** Schematic of the setup for cortical optogenetic inhibition experiments (left), and a schematic of the whole-trial inhibition (right), lasting from 200 ms before the tone or stimulus until the end of the 500-ms response window, followed by a 200-ms ramp-down period. Illumination was performed through the intact skull by a blue laser coupled to X- and Y-scanning galvanometric mirrors. **(b)** Light spots were projected onto each of 42 locations aligned to the Allen Mouse Common Coordinate Framework, both ipsilaterally and contralaterally to the whisker stimulus, with the ∼40% inhibition radius (1.5 mm) plotted below. Locations were separated by 1.15 mm horizontally and 1.4 mm vertically, with an additional vertical shift of 0.25 mm in lateral regions to target the center of S1-bc and whisker region of S2. **(c)** Effects of laser inhibition in each region on the lick rate across all stimulus trials in standard (left) or reversed (right) blocks. Circle size indicates the significance of the effect while its color indicates the effect direction and size. **(d)** Simulations of a decrease in sensitivity (top row) or in lick bias (bottom row) in standard blocks alone (left-hand column) or the full blocks task (right-hand column). **(e)** Effects of laser inhibition in each region on the stimulus strength at curve intersection (i.e., horizontal shifts, top), and on the lick rate at curve intersection (i.e., vertical shifts, bottom). Circle size indicates the significance of the effect (same as in c) while its color indicates the effect direction and size. **(f)** Psychometric effects of inhibition in a subset of regions. From left to right: contra. S1-bc (contralateral barrel cortex), contra. S2 (contralateral secondary somatosensory cortex), ipsi. S1-bc (ipsilateral barrel cortex), contra. M2 (contralateral premotor cortex), ipsi. M2 (ipsilateral premotor cortex), bi. M2 (bilateral premotor cortex), bi. PPC-A (bilateral posterior parietal cortex, area A), and bi. RSC (bilateral retrosplenial cortex). Significance of shifts was tested by comparison with shuffled data and corrected for multiple comparisons using the Dunn–Šidák correction. Arrows are displayed for shifts with P < 0.05. * *P <* 0.05, ** *P <* 0.01, *** *P <* 0.001. Mouse illustration in (a) was partially created with BioRender.com.

A change in lick rate could be caused by an effect on sensitivity to the vibrotactile stimulus, which corresponds to the animal’s perceptual judgments, or an effect on lick bias, which corresponds to the animal’s motivation or ability to lick. To examine how we can distinguish between the two, we performed psychophysical simulations using our logistic classification model and compared the resulting fit parameters and false alarm rates to the ground truth (example in Fig. 2d, left; schematic in Extended Data Fig. 3a). We found that both qualitatively and quantitatively, decreases in sensitivity and lick bias may be difficult to distinguish even with a high number of trials (1000 trials per condition; Extended Data Fig. 3c). However, when considering the psychometric curves of both blocks in conjunction (Fig. 2d, right), the former resulted in a horizontal shift of the intersection of the curves, as mice licked less in response to the stimulus during standard blocks but more during reversed blocks, while the latter resulted in a vertical shift, as mice licked less during both blocks. Simulations confirmed that intersection shifts better reflected underlying changes in sensitivity and bias than either fit parameters (i.e., bias and sensitivity) or false alarm rates (Extended Data Fig. 3b–e).

We found that horizontal shifts were localized to the contralateral S1-bc and M2 (Fig. 2e; detailed effects in Extended Data Fig. 2e). We also observed small vertical shifts that had a bilateral pattern across hemispheres but were slightly stronger on the ipsilateral side, such as a negative shift upon inhibition of either barrel cortex and a positive shift upon inhibition of either medial motor cortex. To more precisely characterize the effects of inhibition, we repeated the experiment in a subset of regions with seven stimulus intensities (Fig. 2f; *n* = 31,402 trials across 5 mice). For unilateral M2 inhibition, we selected a location in the center of the four effective spots from the optogenetic screen, which corresponded to the anterolateral motor cortex (ALM). Note, however, that the measured extent of inhibition means it extends considerably beyond the functional radius of the ALM, which was estimated at 0.75 mm^46^. In addition, we examined the effect of bilateral inhibition of M2 and two associative cortical regions where we did not find a unilateral effect, but which were previously found to be involved in certain types of perceptual judgments—area A of the posterior parietal cortex (PPC-A)^3,47^ and retrosplenial cortex (RSC)^13,14^. Bilateral M2 inhibition caused the largest horizontal shift observed. The results for unilateral somatosensory and unilateral M2 inhibition agreed with the cortex-wide screen, whereas bilateral inhibition of PPC-A caused only a small horizontal shift and bilateral inhibition of RSC with four inhibition spots (to fully cover this elongated structure) had no apparent effect (Fig. 2f). None of the inhibitions significantly increased air licks (Extended Data Fig. 4a), but they affected lick latencies in a manner consistent with the effects on lick rates (Extended Data Fig. 4b).

While previous studies have implicated M2 in high-level sensory processing beyond motor planning and execution^48,49,19,50,51^, it is nonetheless surprising that it plays a role in detection, which is commonly considered a lower-level process. Moreover, this role was apparently dominant over any motor role as bilateral M2 inhibition *increased* licking after stimuli in the reversed blocks. To verify this effect could generalize beyond the blocks task, we repeated the experiment in a delayed lick-left/lick-right detection task (Extended Data Fig. 5a-d). Here, mice were trained to lick either to the same side (congruent contingency) or the opposite side (incongruent contingency) of the same left whisker stimulus (*n* = 3 mice in each contingency, of which two mice were trained in both). Three features distinguished this task from ours: (1) the mice chose between licking left vs. right, as opposed to licking left/right vs. not licking, (2) there was a 600 ms delay between the beginning of the whisker stimulus and the response cue, during which we delivered optogenetic inhibition, and (3) mice did not switch contingencies in the same session. In psychophysical simulations of this task, we found that the shifts approach across contingencies again distinguished an effect on sensitivity vs. an effect on bias, where this time the bias was of licking towards one of the two sides (Extended Data Fig. 5e). Applying this approach to the resulting data, we found that contralateral S1-bc inhibition decreased sensitivity, albeit to a lower extent than in the blocks task and without affecting bias (Extended Data Fig. 5f). Ipsilateral M2 inhibition caused a similarly small impairment in sensitivity alongside a pronounced ipsilateral (left) lick bias, in agreement with previous studies^18,52^. In contrast, either contralateral or bilateral M2 inhibition strongly reduced sensitivity, causing mice to lick more toward the no-stimulus side regardless of whether it was on the left or right. Our findings thus show that the sensitivity impairment we observed when inhibiting M2 was general to vibrotactile detection rather than being due to any specific feature of the blocks task.

### Relating cortex-wide perceptual judgment coding to action coding

Our optogenetic findings indicate that not only the somatosensory cortex but also the premotor cortex is causally important for perceptual judgments as opposed to actions. This suggests that activity in sensory regions alone does not determine perceptual judgments. However, we also wanted to know how perceptual judgments are encoded, and whether this encoding is independent of the resulting actions. To address this question cortex-wide, we performed widefield imaging in transgenic mice expressing GCaMP6s in excitatory cells^53^. Fluorescence was recorded through the cleared, intact skull and aligned to the Allen Mouse Common Coordinate Framework (CCF v3; Fig. 3a). Previous studies have shown that widefield calcium signals reflect combined somatic and neuropil activity primarily from layers 1 to 3^54,55^.

**Figure 3:**
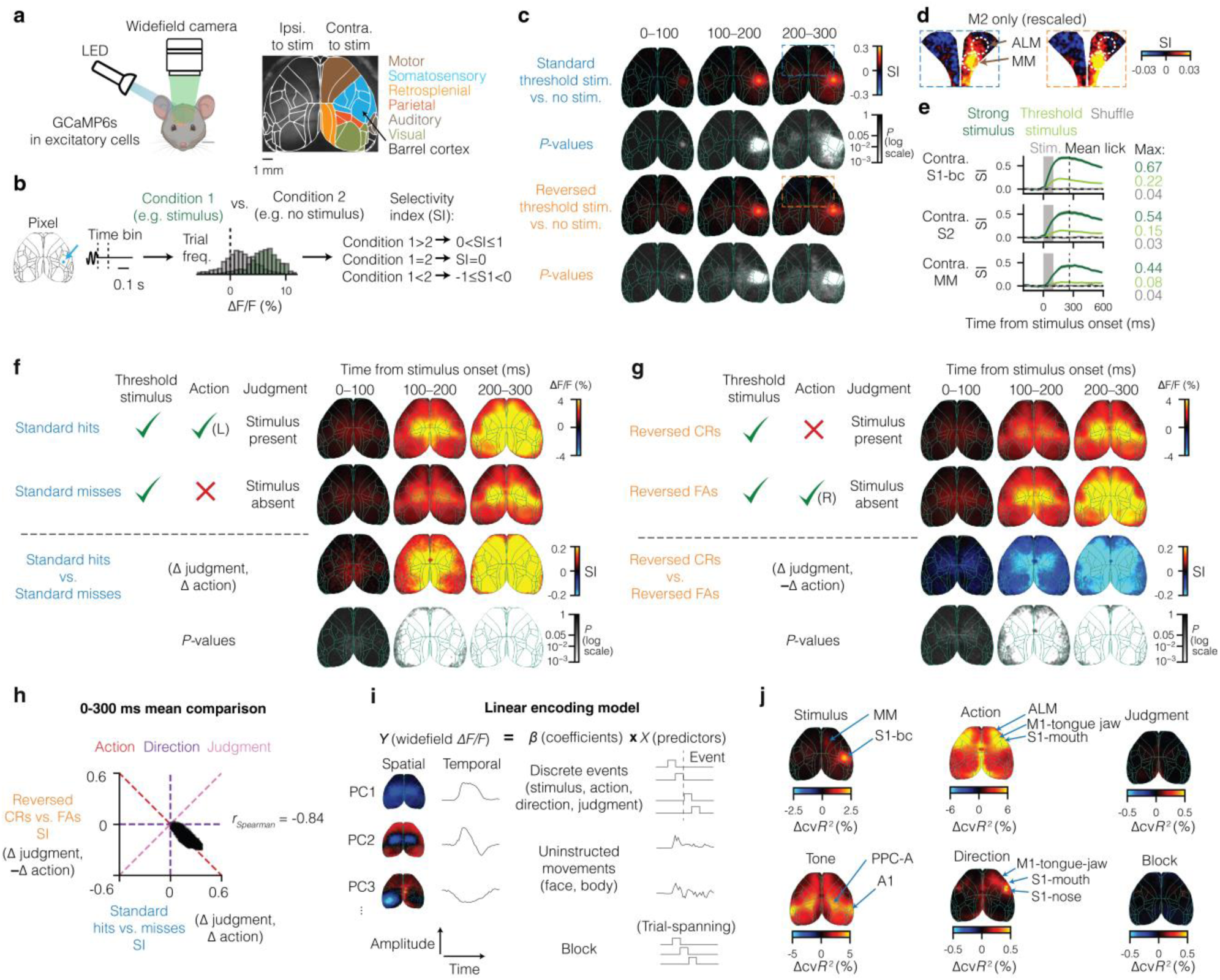
Cortex-wide correlates of perceptual judgments are coupled to actions. **(a)** Widefield calcium recording setup (left), and alignment of widefield imaging to Allen Institute CCF v3, labeled by region category within the hemisphere contralateral to the stimulus (right). **(b)** Selectivity index (SI) calculation. For each pixel and time bin (left), we extracted the distributions of single-trial responses from different trial types (right), and then calculated SI as a normalized measure of the distance between the distributions. **(c)** Results of SI analysis comparing all trials with a threshold whisker stimulus to all trials without a whisker stimulus, separately for the standard (top two rows) and reversed contingency (bottom two rows) blocks. For each comparison, we display the color-coded magnitudes of the SIs and the Benjamini–Hochberg-corrected *P* values of the SIs obtained by comparing them to shuffled results (Methods). All frames within each time bin were averaged for this calculation. **(d)** Rescaled SI results for threshold stimulus vs. no-stimulus trials in M2 alone, with ALM (anterolateral motor cortex) and MM (medial motor cortex). **(e)** Comparison of block-averaged stimulus response time courses in contra. S1-bc (contralateral barrel cortex), contra. S2 (contralateral secondary somatosensory cortex), and contra. MM (contralateral medial motor cortex) across threshold and strong stimulus intensities as well as a trial shuffle (chance-level SI). Error bands show mean ± s.e.m. across *n* = 7 mice. **(f)** Comparison of stimulus present vs. absent judgment trials in the standard blocks. From top to bottom: grand-averaged activity in standard hits, grand-averaged activity in standard misses, the results of the SI calculation between the two trial types, and the Benjamini–Hochberg-corrected *P* values of the SIs obtained by comparing them to shuffled results. The columns on the left represent the three factors defining each trial type (stimulus, action, and judgment), while the columns on the right represent the three time bins from stimulus onset. **(g)** Same as (f) but for stimulus present vs. absent judgment trials in reversed blocks. **(h)** Scatter plot of all widefield pixels across the comparisons in (f) and (g) revealed a strongly negative relationship on a pixel-wise basis. Labels at the top are colored like the axes and diagonals along which action-coupled, direction-coupled, and action-independent perceptual judgment correlates are expected to reside. **(i) S**chematic of the linear model. We aimed to predict widefield activity (left) from the product of coefficients and predictors (right). Widefield activity and uninstructed movements were both decomposed into principal components. **(j)** Unique explained variances corresponding to each predictor, with labeling of peak regions. Note the differences in scale between the predictors. Error bands show mean ± s.e.m. across *n* = 7 mice. Mouse illustration in (a) was partially created with BioRender.com.

We first asked what cortical regions responded to the whisker stimulus, and whether this response differed across blocks, by computing the selectivity index (SI) across stimulus and no-stimulus trials in individual pixels and time bins (Fig. 3b; Methods). As expected, we found that the stimulus response was strongest in the barrel cortex (S1-bc) contralateral to the stimulus, where it appeared as a peak at the central C2 barrel (Fig. 3c), with considerable activity spread to the laterally adjacent S2. Additionally, we observed a peak in the medial M2 in a subregion known as the medial motor cortex (MM; Fig. 3d)^46^. This region is distinct from the previously targeted anterolateral motor cortex (ALM), which is located more anteriorly (∼1.4 mm away). Nonetheless, it was still within the radius of the targeted inhibition (Fig. 2f), and was also directly inhibited in our optogenetic screen (region 17, Extended Data Fig. 2e), with both inhibitions decreasing stimulus sensitivity. Responses in S1-bc, S2, and MM were similar across blocks (Fig. 3c,d), were strongly dependent on stimulus intensity (Fig. 3e), and showed trial-to-trial variability that was correlated across regions (Extended Data Fig. 6a,b).

Given that our stimulus response calculation included both stimulus present and absent judgments, we wondered if the observed trial-to-trial variability reflected the perceptual judgments. To test this, we compared trials in which the same threshold stimulus was given, but in which the mouse made either a stimulus present or absent judgment. In the standard blocks, we compared hits (mouse received a whisker stimulus and licked, meaning it made a stimulus present judgment) to misses (mouse received a whisker stimulus and did not lick, meaning it made a stimulus absent judgment). Both displayed a whisker response in S1-bc, together with a bilateral response in medial regions. The latter appeared to be related to uninstructed movements caused by the tone^56^ (i.e., startle response; Extended Data Fig. 6c) as it was observed across all trial types, including no-stimulus, no-lick trials^56^ (Extended Data Fig. 6d; Supplementary Fig. 2). Overall, however, activity was greater in standard hits, particularly around medial regions but also elsewhere throughout the cortex (Fig. 3f).

In the reversed blocks, we compared correct rejections (CRs; mouse received a whisker stimulus and did not lick, meaning it made a stimulus present judgment) to false alarms (FAs; mouse received a whisker stimulus and licked, meaning it made a stimulus absent judgment). If the differences between hits and misses in the standard block reflect the perceptual judgments in an action-independent manner, similar positive differences would be expected in the reversed blocks. However, the differences were significantly negative throughout the cortex, consistent with general (i.e., lick vs. no lick) action coding (Fig. 3g). Moreover, they were strongly inversely correlated across blocks, indicating they were coupled to action (Fig. 3h), rather than reflecting judgment in an action-independent manner. This was also true when considering only trials in which mice licked later than 300 ms from stimulus onset and quantifying the evolution of activity using the finest possible temporal resolution of 30 Hz (Extended Data Fig. 6e).

It is possible that the action-independent correlates of perceptual judgments were simply too weak to detect due to being overshadowed by the much larger action correlates in the same cortical regions. To test this, we applied ridge regression^35^, which modeled widefield activity as a linear sum of predictors and allowed us to tease apart the influence of each predictor (Fig. 3i). These predictors included discrete events (stimulus, action, direction, judgment), uninstructed movements (face and body), and the block context. For event predictors, we also added time-shifted copies to account for neural correlates before or after each event (Methods). Importantly, Singular value decomposition revealed that 99% of the variability in widefield data was contained in 113 ± 9 principal components (mean ± s.e.m. across 7 mice; Extended Data Fig. 6f), which our model aimed to predict. We calculated the best-fitting coefficients for each predictor and performed cross-validation to assess the model’s predictive power (cross-validated explained variance, cv*R²*). To conservatively estimate each predictor’s contribution, we fitted reduced models by excluding each predictor and calculated the reduction in cross-validated explained variance (unique explained variance, Δcv*R*^2^).

Our results for the unique explained variance of the stimulus replicated the earlier findings (Fig. 3j). Meanwhile, the unique explained variance for the action predictor was strongest in lateral regions, particularly the mouth somatosensory cortex and tongue-jaw M1, rather than medial regions as before. This is likely due to action being correlated with increased uninstructed movements in the face and body (Fig. 1f), which accounted for much of the variance in medial regions (Extended Data Fig. 6g). Tone-evoked activity was nearly global but strongest in A1 and PPC-A, as previously observed^57^. The direction predictor explained modest activity in the right mouth and nose somatosensory cortices and tongue-jaw M1, similar to the action predictor but more strongly in the right hemisphere. However, the perceptual judgment predictor (the main predictor we aimed to isolate with this approach) and block context predictors did not appreciably explain activity anywhere on the cortical surface, even when averaged across all pixels in key regions from the optogenetic screen or the entire dorsal cortical surface (Extended Data Fig. 6h). Thus, the regression analysis supports our previous results in finding that perceptual judgments are not represented independently of actions in cortical activity.

Lastly, we tested whether our findings generalize to the delayed lick-left/lick-right task as before. In the congruent contingency (Extended Data Fig. 7a), we compared hits (mouse received a whisker stimulus and licked left, meaning it made a stimulus present judgment) to misses (mouse received a whisker stimulus and licked right, meaning it made a stimulus absent judgment). In the incongruent contingency (Extended Data Fig. 7b), we similarly compared hits and misses, but now the lick directions in each trial type were reversed. Surprisingly, in both contingencies we found a significant positive difference beginning at about 200 ms post-stimulus (400 ms before the response tone), which had a similar bilateral spread to that in standard blocks (Fig. 3f) but with a lower magnitude. This resulted in the hit vs. miss difference being significantly positively correlated across contingencies (Extended Data Fig. 7c). While this result appears to imply widespread action-independent judgment coding in this task, we also noticed that mice made more uninstructed movements during the delay in hits compared to misses in both contingencies (Extended Data Fig. 7d). This contrasted with the reversal observed across contingencies in the blocks task (Fig. 1f) and suggests that the observed hit vs. miss differences may reflect uninstructed movements rather than judgments per se. Indeed, three additional results confirmed this: (1) pairwise differences across trials in both contingencies were correlated to differences in uninstructed movements (Extended Data Fig. 7e), (2) subselecting hits and misses to equalize the magnitude of uninstructed movements (i.e., taking the ∼50% of hit trials with less movement and ∼50% of miss trials with more movement) eliminated the positive difference between them (Extended Data Fig. 7f,g), and (3) a linear regression model showed that judgments do not account for additional variance in neural activity beyond that which uninstructed movements account for (Extended Data Fig. 7h). In sum, it appears that during an imposed delay in the lick-left/lick-right task, judgment correlates become coupled to *uninstructed* movements rather than instructed ones as in the blocks task.

### Relating single-cell perceptual judgment coding to action coding

As widefield imaging captures the aggregate activity of many cells at each location, we wondered if examining single-cell signals could provide additional insights. For example, perceptual judgments might be encoded sparsely^4^ or across orthogonal dimensions in the same neural population^58,59^, which could render them invisible to widefield imaging. To investigate this, we recorded two-photon calcium activity during the blocks task in individual cortical layer 2–4 excitatory cells in regions that we believe play a causal role in perceptual judgments based on the previous findings: S1-C2, MM, and ALM. Additionally, we recorded from PPC-A and RSC, despite their small or insignificant causal role (Fig. 2), due to these regions’ association with perceptual judgments^3,13,14,47^. Data were collected from 2–3 fields of view (FOVs) per mouse and region, with 335 ± 135 cells per FOV (mean ± s.d.; Fig. 4a–c). Activity was normalized to the pre-trial baseline (*ΔF/F*) as before. Cells showed diverse task-related responses, including encoding of stimulus, action, and direction (Fig. 4d–f; Supplementary Fig. 3).

**Figure 4:**
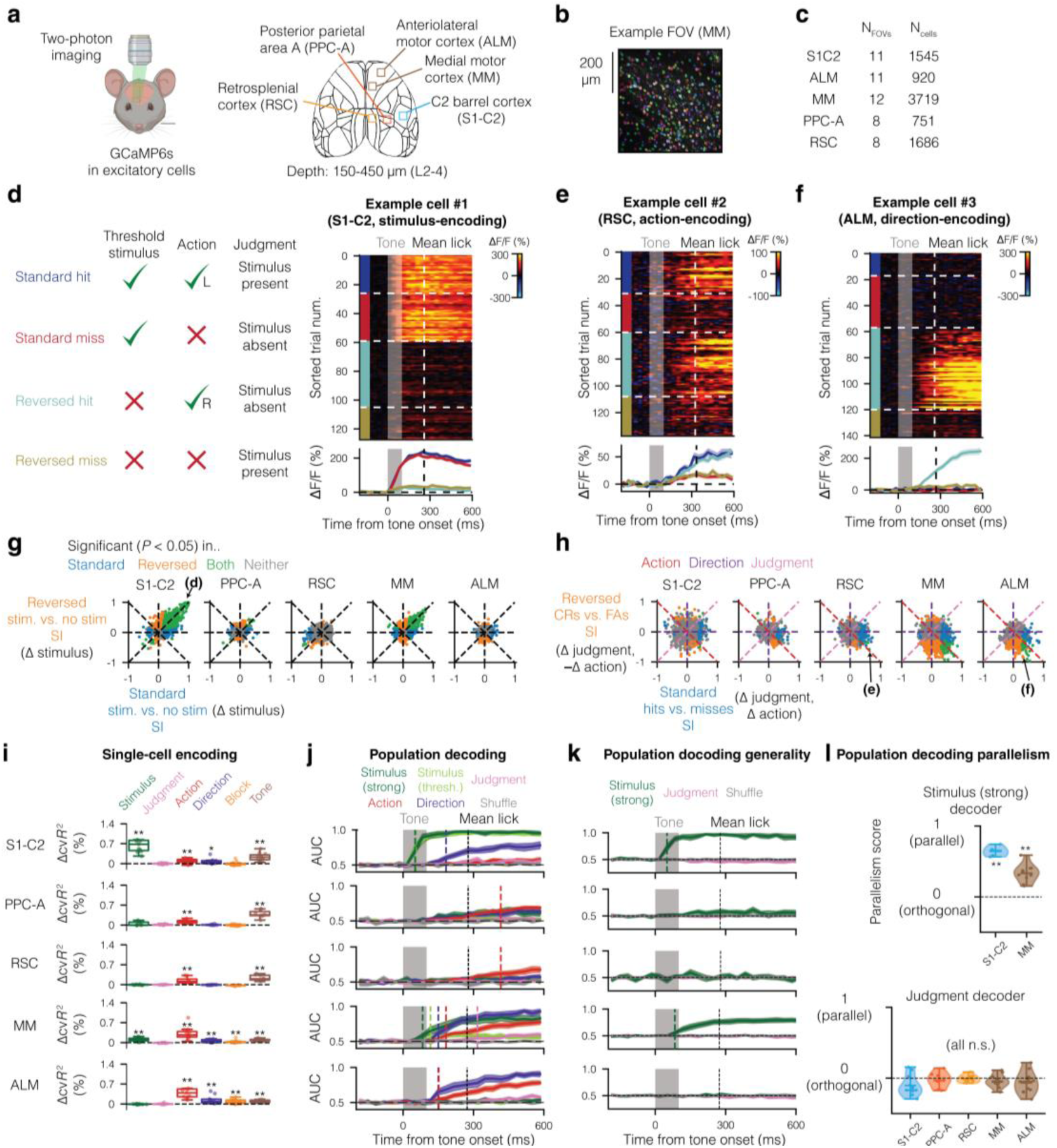
Single-cell correlates of perceptual judgments are coupled to actions. **(a)** Two-photon imaging of CaMKII-GCaMP6s mice under a 20× objective (left) and the imaged regions (right). **(b)** An example field of view (FOV) from the medial motor cortex (MM), with detected cells colored randomly. **(c)** Number of fields of view and cells recorded in total from each cortical region. **(d)** An example of an S1-C2 stimulus coding cell, with trial labels on the left, raster plot on the top-right, and peri-stimulus time histogram (PSTH) on the bottom-right. This cell selectively responds during stimulus trials, regardless of action and judgment. **(e)** As in (d), but for an RSC action coding cell. **(f)** As in (d), but for an ALM direction coding cell. **(g)** Comparison of stimulus coding in the standard vs. reversed-contingency blocks shows the two are positively correlated in stimulus-coding regions. Cells with a significant SI (*P <* 0.05 after correction for multiple comparisons using the Benjamini–Hochberg procedure) from only the comparison in the standard block, only the comparison in the reversed block, both, or neither are plotted in blue, magenta, green, and gray, respectively. **(h)** Comparison of SIs for reversed CRs vs. FAs and for standard hits vs. misses. Point colors as in (g). Labels at the top are colored like the axes and diagonals along which action-coupled, direction-coupled, and action-independent perceptual judgment correlates are expected to reside. **(i)** Unique explained variances corresponding to each predictor, averaged across all cells in each session/field of view. Boxes correspond to the 1st to 3rd quartiles, with the line representing the median and whiskers extending to the farthest data point within 1.5× the interquartile range (IQR). Dots correspond to individual sessions/fields of view. Stars above each predictor indicate its significance relative to zero (one-sided Wilcoxon signed-rank test, corrected for multiple comparisons using the Benjamini–Hochberg procedure). **(j)** Performance of cross-validated logistic classifiers with L1 regularization, run separately across regions (subpanels), factors (lines, see legend on top), and time after the tone (ordinate). Colored vertical dotted lines represent the first time at which the respectively colored factor decoding first reached significance (*P* < 0.05, two-sided Wilcoxon signed-rank test, FDR-corrected). Darker lines show means with shading representing s.e.m. across fields of view. **(k)** Results of cross-block population decoding. Trial types were split into two balanced sets for training and testing (Methods). The AUCs shown are the mean AUCs of first-set-to-second-set and second-set-to-first-set decoders for each time point. Colored vertical dotted lines represent the first time at which the respectively colored factor decoding first reached significance. **(l)** Parallelism score across sets for the strong stimulus decoder (top, only regions with significant decoding included) and judgment decoder (bottom, all regions included). Stars above each region indicate the significance of the score relative to zero (one-sided Wilcoxon signed-rank test, corrected for multiple comparisons using the Benjamini–Hochberg procedure). Error bands in (d), (e), (f), (j), and (k) are s.e.m. across trials in each session. * *P <* 0.05, ** *P <* 0.01, *** *P <* 0.001. Mouse illustration in (a) was partially created with BioRender.com.

We started by comparing the selectivity index (SI) for the whisker stimuli, regardless of the action, across the standard and reversed blocks, as in the widefield analysis but now in single cells. SIs were both significant and highly correlated across blocks in S1-C2 and MM, consistent with the widefield results (Fig. 4g), suggesting that in these two regions many cells encoded the stimulus. Next, we examined the SIs across perceptual judgments in the standard and reversed blocks, respectively computing them from standard hits vs. misses and reversed CRs vs. FAs (Fig. 4h). Again, we reasoned that cells encoding action-independent judgments would significantly encode the judgment across both comparisons, putting them on the identity line. However, only a handful of cells satisfied this criterion after correcting for false discovery rate (3/1545 cells in S1-C2, 1/3719 cells in MM, and none elsewhere). The same was also true when restricting the analysis to trials in which the mouse licked after 300 ms, and examining the evolution of SIs on a frame-by-frame basis (Extended Data Fig. 8a). To see how perceptual judgments are encoded in the cells in which this coding was significant, we examined the raster plots and PSTHs of three of the highest-SI cells across both comparisons (Extended Data Fig. 8b-d). Coding was weak in each case, with large trial-to-trial variability resulting in small and nearly overlapping average differences between the trial types. Many more cells in Fig. 4h were significant in only the standard (blue dots) or reversed (magenta dots) blocks, particularly in MM and ALM, suggesting they encoded the lick direction (left or right lick). Still others were on the anti-diagonal, suggesting they encoded the lick action. Thus, in contrast to the widefield perceptual judgment coding (Fig. 3h), which was largely coupled to the lick action, single-cell perceptual judgment coding was additionally strongly coupled to the lick direction. To systematically assess which factors could be combinatorially influencing cell activity, we built a ridge regression model as before, but now at the single-cell level. We then quantified the unique variance explained (Δc*vR²*) by each predictor, averaged across cells in each region (Fig. 4i). Consistent with prior analyses, the stimulus factor was significant only in S1-C2 and MM, and the perceptual judgment factor was not significant anywhere. Likewise, although smaller in magnitude, action and tone correlates were significant cortex-wide. However, direction and block coding emerged in regions where they did not appear in widefield data, namely contralateral MM and S1-C2 for the former and contralateral MM and ALM for the latter.

While individual cells did not strongly encode perceptual judgments independently of actions, it is possible that in conjunction, several weakly encoding cells could allow a downstream decoder to read out the judgment. To test this, we used cross-validated logistic classification on all simultaneously recorded cells in each FOV (Fig. 4j; Methods). Performance was measured with AUC (area under the receiver operating characteristic curve), which ranges from chance level at 0.5 to perfect at 1. The classifier nearly perfectly identified threshold and strong stimuli in S1-C2 (max AUC > 0.95) and was well above chance (max AUC = 0.83) for strong stimuli in MM, consistent with the widefield observations (Fig. 3e). It also classified action fairly well, reaching an AUC of 0.82 in the ALM. However, classification of perceptual judgments across both blocks was poor (max AUC ≈ 0.6), only slightly exceeding chance level after the mean lick onset in MM and ALM (>300 ms). These results held across a variety of classifiers, including non-linear ensembles (Supplementary Fig. 4). Next, as a more stringent test of perceptual judgment coding, we examined to what extent this coding generalized across different trial types^60^. To do so, we trained a judgment decoder on one block-, stimulus-, and action-matched set of trial types and tested it on another set of matched trial types, with a similar procedure applied to strong stimulus coding as a control (Methods). While stimulus coding generalized in S1-C2 and MM as expected, judgment decoding failed to generalize in any region, yielding shuffle-like performance (Fig. 4k). Lastly, we computed the parallelism score (PS)^60^ across the same splits (Methods). PS is a measure of the extent to which two decoder weight vectors are parallel, indicating consistency in the population coding. We found that strong stimulus coding was significantly parallel across splits in both S1-C2 and MM, while the judgment coding was orthogonal in all regions (Fig. 4l). Overall, our findings indicate that stimulus presence judgments are coupled to actions rather than being encoded independently in the recorded regions, on both the single-cell and population levels.

### Block coding in premotor regions

Our findings so far indicate that the premotor cortex is causally important for stimulus sensitivity, but premotor cortical activity is well accounted for by the actions themselves. While for simple behaviors (e.g., reflexes), direct activation of action by stimulus inputs seems sufficient, it is unclear how mice were able to switch between performing different actions in response to the same vibrotactile stimulus (or lack thereof) in our task. At the same time, the regression model revealed significant coding of the block in MM and ALM (Fig. 4i). Because this coding is potentially of interest in how mice solved our context-based task, we wanted to explore it in more depth. We began by examining block coding in individual cells by calculating each cell’s block selectivity index across no-stimulus, no-lick trials in each block (Methods). After correcting for multiple comparisons (population false discovery rate), we found that 10.6 ± 3.7% and 9.8 ± 1.4% (mean ± s.e.m. across mice) of cells in MM and ALM, respectively, significantly encoded the block relative to shuffled data.

Two block-coding cells from a single ALM recording session are plotted in Fig. 5a. The two reliably alternated in activity according to the block. Moreover, their activity was locked to the tone onset, regardless of stimulus or any subsequent licking (Fig. 5b). To test whether the block context correlates were tone-locked in general, we ran a logistic block classifier using all simultaneously recorded cells across no-stimulus, no-lick trials. Indeed, the cross-validated performance of the classifier departed from chance level in MM and ALM shortly after the tone ended and reached an AUC of 0.8, but remained near chance level in the other regions (Fig. 5c). We confirmed these results using deconvolved activity, which was not baseline-corrected (Extended Data Fig. 9a). In addition, we repeated the analysis after excluding the significant block cells, which reduced classifier accuracy to near chance levels (Extended Data Fig. 9b), and using only significant block cells, which even slightly improved accuracy compared to using the full population (Extended Data Fig. 9c). In sum, it appears that these regions do not maintain a persistent representation of the block context but rather transiently represent it following the tone, relying on a small subpopulation of block-encoding cells.

**Figure 5:**
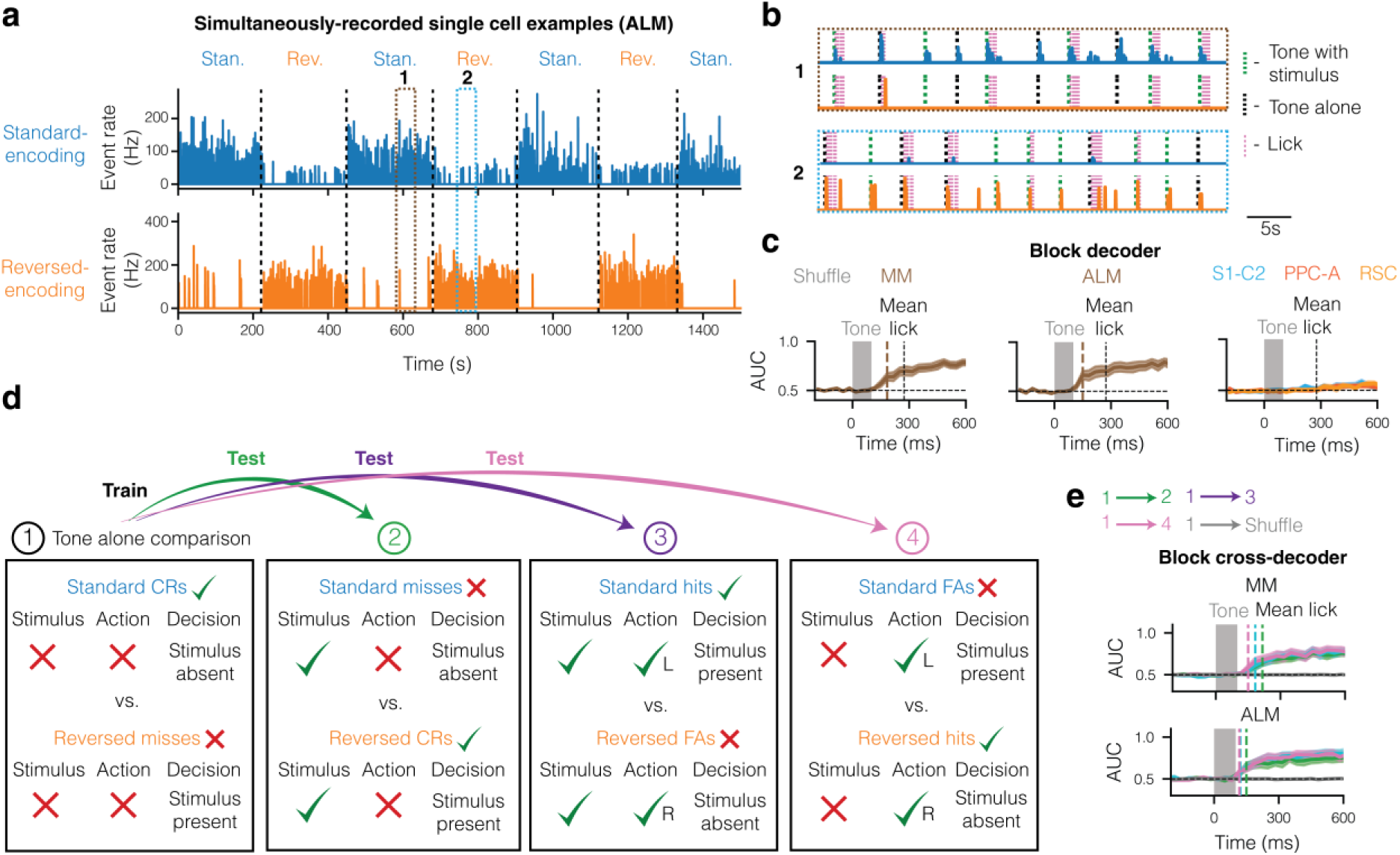
Transient block coding in premotor regions. **(a)** Deconvolved event rates of two cells recorded simultaneously from layer 2 of the ALM in one session, namely a standard block cell (top) and a reversed block cell (bottom). **(b)** Zoom-in on the numbered sections in (a) shows consistent standard block cell activity following the tone during a standard block (1, top) and weak reversed block cell activity in the same trials (1, bottom), and vice versa during a reversed block (2, top and bottom). **(c)** Performance of cross-validated logistic classifiers of the block using activity from MM (left), ALM (center), and other regions (S1-C2, PPC-A, RSC; right). Colored vertical dotted lines represent the first time at which the block factor decoding first reached significance (*P* < 0.05, two-sided Wilcoxon signed-rank test, FDR-corrected). **(d)** Trial type pairs on which the block cross-decoder was trained (1, leftmost box) or tested (either 2, 3, or 4). Checks or crosses next to trial type names indicate the correctness of the mouse’s behavior. **(e)** Results of the cross-decoder for each test in MM (top) and ALM (bottom). Error bands in (c) and (e) are s.e.m. across sessions in each region.

Does block coding depend on the particular trial type? We examined this by training a classifier to decode the block using all recorded cells across tone-only trials (standard CRs and reversed misses), which differed in block, perceptual judgment, and correctness, and testing it on other trial-type comparisons across blocks in which perceptual judgment, correctness, or both were inverted (Fig. 5d). The results of classification were unaffected by perceptual judgment and correctness, as well as the presence/absence of a stimulus and differences in lick direction (Fig. 5e). This was also the case when examining the mean response magnitudes of block cells, except for slightly increased responses in lick trials (Extended Data Fig. 9d). Thus, block coding appears to be largely independent of trial type. Additionally, we tested the block classification accuracy on the first trials following block switches (Extended Data Fig. 9e). We did not observe a significant drop in accuracy, suggesting that block coding switched rapidly, consistent with behavioral observations (Extended Data Fig. 1c).

Lastly, given that each block is associated with a different lick direction, we wondered if there is a relationship between the activities of block and direction cells in our recordings. First, we quantified the percentage of direction cells by computing their directional selectivity index and removing overlapping block cells (Methods), finding that 8.9 ± 3.1% and 17.9 ± 8.4% of the cells in MM and ALM, respectively, encoded lick direction but not block. Next, we computed the noise correlation between the two groups of cells at different time lags relative to each other^61^. These correlations quantify shared variance across cells that is not explained by the trial structure, as would be expected if the two were directly connected, although they do not prove connectivity. We observed positive correlations between standard block and lick left cells, as well as between reversed block and lick right cells, consistent with these cell classes being positively connected (Extended Data Fig. 9f). In contrast, the negative correlations between standard and reversed block cells, and between lick left and lick right cells, were consistent with these cell classes being negatively connected, i.e., in competition with each other.

### An action-coupled model of perceptual judgments

The finding of robust block coding in the premotor cortex, as well as its shared variance with direction coding, offers an interesting hint as to how mice can perform our context-based task. We sought to incorporate this finding into a model of perceptual judgments based on the observed functional cell types, which also included stimulus inputs and direction output. This model consisted of a neural network in which two populations of 100 stimulus and block units, together with noise, synapsed onto two populations of 100 direction units that competed with each other (Fig. 6a), yielding a continuous decision variable (DV) defined as the left–right difference in accumulated inputs. The current block determined the weight of the stimulus inputs (sensitivity), as well as the magnitude and direction of block inputs (akin to bias). These inputs were present during the 100 ms stimulus/tone phase, and then the population of direction cells was allowed to drift (noise only) during the remaining response window. If either the lick left or lick right population reached its respective bound, then the mouse licked in that direction after a fixed time delay given by the non-decision time (NDT). In addition, we added non-direction-selective action feedback coding in accordance with the observed widespread encoding of action (Figs. 3g and 4i,j). Importantly, perceptual judgments were not an explicit component of this model. Rather, they were inferred from the specific action performed in the context of each block (right-hand column in Fig. 6a), making this an action-coupled model.

**Figure 6:**
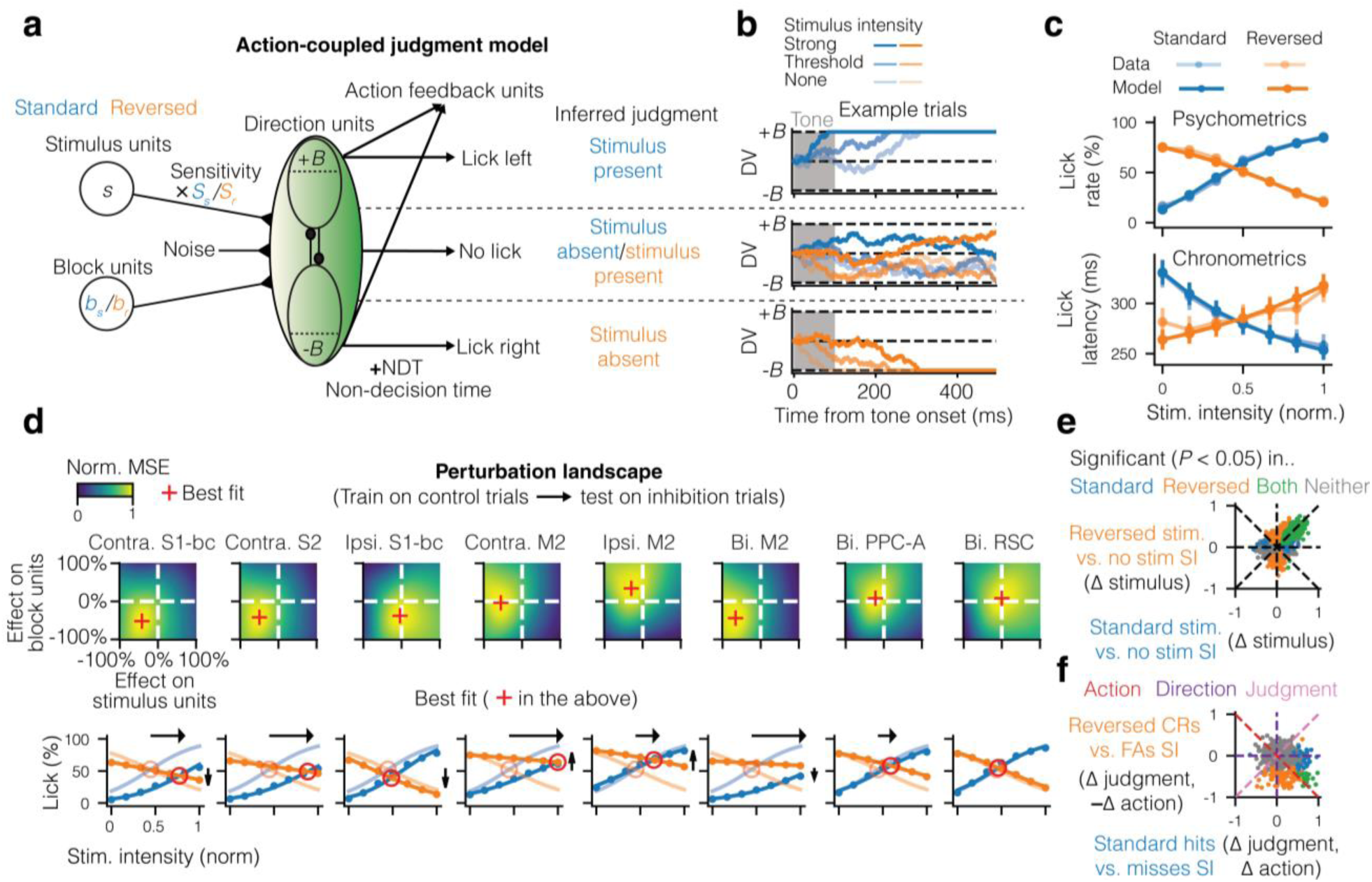
An action-coupled perceptual judgment model parsimoniously accounts for behavior. **(a)** Schematic of the action-coupled perceptual judgment model. The inputs to this model are stimulus units (weighted by sensitivity), block units, and noise, which are integrated to a higher bound, a lower bound, or neither within the response window. Each action is interpreted as a perceptual judgment according to the block, but the perceptual judgments are not represented explicitly. **(b)** Single-trial examples of each of the outcomes in (a), from top to bottom: lick left, no lick, and lick right. **(c)** Fits of psychometric curves (top) and chronometric curves (bottom). **(d)** Comparisons of models trained on control trials with perturbations to the sensitivity (*y*-axis) or block (bias) inputs (*x*-axis) to those obtained from inhibiting each region (top row), and resulting peak fits compared to the data (bottom row). **(e)** Comparison of stimulus coding in the standard vs. reversed-contingency blocks in the model, as in Fig. 4g. **(f)** Comparison of SIs for reversed CRs vs. FAs and for standard hits vs. misses, as in Fig. 4h. Error bars in (c) are s.e.m. across mice. * *P <* 0.05, ** *P <* 0.01, *** *P <* 0.001.

We separately fit the model parameters to reproduce the psychometric and chronometric curves for each mouse (Extended Data Fig. 10a; Methods). Overall, we found that in standard blocks, both stimulus and block inputs were needed to reach the upper (lick left) bound, and in reversed blocks, the block inputs were strong enough on their own to reach the lower (lick right) bound while the stimulus inputs biased the mouse against licking (Extended Data Fig. 10a). The asymmetry of the total drive—that is, how much the combination of stimulus and block inputs pushed the decision variable towards one side in particular—was associated with both a higher lick rate and a lower lick latency (Fig. 6b), which allowed the model to explain the reversed trends across the psychometric and chronometric curves (Fig. 6c).

Next, we tested whether this model could account for the effects of optogenetic inhibition (Fig. 2f). We trained it on the psychometric curve derived from control trials and performed parameter perturbations to examine which perturbations best fit the psychometric effects of inhibition (Fig. 6d; Methods). The resulting best fits closely matched the real psychometric data, with perturbations of block inputs being akin to vertical shifts and perturbations of sensitivity being akin to horizontal shifts. They also reproduced the lick latency effects observed in the real data (Extended Data Fig. 10b). Finally, we performed the same comparisons across trial types as for the measured single-cell activities (Fig. 4g,h), and found that the results qualitatively matched the data (Fig. 6e,f).

Could alternate models also explain behavior in our task? We considered two possibilities. In the first, we optimized a common bias term for both blocks instead of explicit block coding (Extended Data Fig. 10c). As mice could not effectively switch between blocks in this case, it resulted in poor fits to both psychometric and chronometric curves (Extended Data Fig. 10d). In the second, we allowed block coding but only after an initial single-boundary stage that was based on stimulus and bias alone. This was an action-independent perceptual judgment model as crossing the boundary always yielded ‘stimulus present’ judgments while failing to cross it yielded ‘stimulus absent’ judgments, which then mapped to different actions depending on the block (Extended Data Fig. 10e). While this model was able to reproduce the psychometric curves, it could not reproduce the negative slope observed in the reversed block chronometric curve (Extended Data Fig. 10f). This is because licks in reversed blocks corresponded to failures to cross the boundary and therefore could not become systematically faster with stronger stimuli. In addition, this model predicted a class of action-independent perceptual judgment neurons which we did not observe in the cortex. In sum, the action-coupled model, in which block inputs are directly combined with stimulus inputs, parsimoniously accounts for major features of both behavioral and neural data. Although this model is simplistic and unlikely to accurately represent how the brain solves this task, it nonetheless solves it with only the observed functional cell types and without invoking action-independent perceptual judgment coding.

## Discussion

In this study, we set out to determine how sensory stimuli are converted into action using a contingency blocks task that separates perceptual judgments from actions. Based on previous findings, we chose to focus on the neocortex, where we applied unbiased techniques to find which regions causally contribute to or encode the perceptual judgment. The main experimental findings were threefold: (1) the premotor cortex (M2) plays a direct role in stimulus sensitivity, at least as much as somatosensory regions, (2) differences in neural activity observed between stimulus present and stimulus absent judgments are largely explained by instructed and uninstructed movements with no significant contribution from judgments per se, and (3) premotor regions encode the current block in a transient manner following trial onsets. Based on these findings, we created a model that explained perceptual judgments as stemming from competition among action-driving cells that receive stimulus and block inputs. Our findings thus suggest a model of sensorimotor transformations in which stimulus inputs directly drive different actions depending on the context. In addition, they are consistent with an action-coupled view in which perceptual judgments are not independently encoded, but rather emerge from competition between different motor plans^32,33^.

The finding that M2 inhibition selectively affected perceptual judgments rather than actions in our task, indicated by the predominance of horizontal shifts over vertical shifts when inhibiting M2, is in line with studies implicating this region in high-level sensory processing beyond motor planning and execution, including evidence accumulation in discrimination^48,51^, sensory working memory^5,49,62,63^, and top-down sensory predictive processing^64,65^. However, it is still somewhat surprising because detection is often considered a lower-level process. The much larger impact of M2 inhibition on perceptual judgments than on actions suggests that M2 has a more fundamental role in perception than previously realized. Accordingly, previous findings that bilaterally inhibiting M2 in discrimination tasks with two distinct stimuli impaired motor action^3,54,66^ require re-examination, as an impairment in perceptual judgments would result in animals failing to perceive either stimulus and thus withholding a response.

Although cortical inhibition had limited effects on action, action coding was widespread. Here, our study joins similar findings of widespread action coding^4,35,67^, despite its apparent lack of an immediate causal role^19^. Our contribution is that perceptual judgment coding is coupled to this widespread motor activity rather than being independent of it. Notably, the nature of the judgment-action coupling depends on the recording method: widefield imaging showed predominant coupling to action, namely licking vs. not licking (Fig. 3h), while two-photon imaging showed predominant coupling to direction, namely licking left vs. right (Fig. 4h). This likely reflects, at least in part, the fact that widefield imaging mixed signals from nearby cells encoding different directions, thereby increasing apparent action coding and reducing direction coding. Importantly, our findings do not exclude the possibility that judgments had distinct correlates across the two blocks. Such correlates would manifest as direction-coupled with our analysis. However, within our framework, they would nonetheless not be action-independent, because judgment would remain embedded in the block-dependent mapping onto the required motor output. By contrast, action-independent judgment correlates would generalize across blocks despite the change in mapping, similarly to stimulus correlates (Fig. 4k,l).

These observations help interpret choice correlates reported in animals performing perceptual tasks. Such correlates are typically defined operationally by the animals’ perceptual report and therefore potentially include activity related to both the perceptual judgment and the expectation, planning, and execution of the report itself^68^. One common scheme is to define choice as action versus inaction (e.g., in a go/no-go task)^3,67,69–71^, while another common scheme is to define it as one action versus another (e.g., in a two-alternative-choice task)^4,9,18,72^. It has been noted that these two report schemes have resulted in different findings, namely that choice coding in the former case is stronger and more widespread^73,74^, and it had been unclear to what extent this difference can be explained by the difference in report scheme versus the perceptual judgment being studied (e.g., detection vs. discrimination). Our findings suggest that the report scheme plays a paramount role in choice correlates and may be sufficient to explain differences across studies.

Especially interesting was our observation of transiently active block cells in the MM and ALM subregions of M2, which can be considered a type of context coding. Although sparse, these cells allowed us to accurately decode the block at the population level (Fig. 5c). Another recent study found that MM and ALM encoded the block in a sensory selection task (lick-to-touch vs. lick-to-light) even before the stimulus onset^75^; notably, in this study, unlike in ours, there was no external indication of the current block on each trial (the lick port side). Combined with the observation that mice rarely performed air licks after block switches (Extended Data Fig. 1c), it is plausible that the transient nature of block coding reflects a sensory-based rather than purely cognitive block identification. This may also explain the counterintuitive finding that inhibiting the somatosensory cortex during the blocks task reduced licking (Fig. 2f), as doing so may have reduced the feedforward somatosensory drive indicating block identity. In contrast, in the delayed lick-left/lick-right task (Extended Data Fig. 5) the contingency was fixed inside sessions, and somatosensory cortex inhibition had no such effect. Additional work is needed to explore how context coding may depend on the availability of external context cues, e.g., by studying both cued and uncued block switches in the same paradigm.

In any case, these block cells provide a task-specific solution to a problem common in perceptual decision-making: how does the brain trigger the appropriate action in response to a stimulus^76,77^? In our model, these cells selectively trigger direction cells, coupling perceptual judgment coding to specific actions based on context. While our model is simplistic, the real brain has a vastly expanded repertoire of solutions available that may also be based on similar principles. For instance, under the view that M2 uses recurrent attractor dynamics to drive action^78–80^, block cells could establish the fixed points for said attractors. This would allow the whisker stimulus to push the system either toward a “lick left” bound in standard blocks or away from a “lick right” bound in reversed blocks, with these boundaries representing the opposite ends of the attractor.

While our results do not definitively answer where the sensorimotor transformation takes place, they suggest some important constraints. First, we found that neither S1 nor S2 inhibition, despite abolishing nearly all activity at the targeted location (Extended Data Fig. 2c), fully abolished stimulus sensitivity. M2 inhibition also did not fully abolish sensitivity, even when the ALM (Fig. 2f) or MM (region 7 in our optogenetic screen; Extended Data Fig. 2e) were specifically targeted. This suggests that the stimulus inputs in our model do not arrive entirely via S1 or M2, but may involve both regions in parallel. They may also arrive via subcortical pathways including the higher-order thalamus^81^, superior colliculus^82^, and cerebellum^83^. Second, although we found both block and direction coding in M2, it is unlikely that all cells of either category were located there. This is because in that case, bilateral M2 inhibition would entirely abolish licking, whereas we only observed a small negative vertical shift (Fig. 2f). We thus suspect that both types of cells were part of a broader cortical-subcortical network, with the accumulation-to-bound process suggested by our model implemented either partially or fully in subcortical regions.

Indeed, the fact we did not record from subcortical regions is an important constraint of this study. While we cannot exclude the presence of action-independent judgment coding in these regions, we think it is unlikely given recent studies that have shown the cortex and subcortex work in tandem and tend to share coding^40,82,84^. In any case, our claims here only concern cortical coding of judgments. Future studies could record or inhibit candidate subcortical regions while animals perform a task that dissociates judgments from actions to examine the subcortical role in sensorimotor transformations.

Lastly, while our results show that perceptual judgment coding is action-coupled in the cortex of mice performing vibrotactile detection, we want to acknowledge that this may not be universal across all perceptual judgments and suggest two possible compromises: one task-dependent and one phylogenetic. The task-dependent compromise posits that while mice may have learned to bypass action-independent perceptual judgment coding in our task over the course of training, such coding would still play a crucial role in more natural and/or complex perceptual judgments. In these cases, circumstances may require that perceptual judgments are abstracted away from any particular action plan. Our findings underscore the need to separate perceptual judgments from actions in other tasks to test for action-independent judgment coding, while also suggesting that such independent representations are not a necessary ingredient in all perceptual behaviors. Meanwhile, the phylogenetic compromise posits that action-independent perceptual judgment coding is prominent in primates but not in mice. Indeed, there is evidence of such coding in non-human primates performing categorization tasks^16,85,86^, and in humans performing even simple detection tasks^87–89^. The possibility that it arises in non-human primates and humans but not in mice aligns with the perspective of Cisek & Kalaska^33^, who argue that the capacity for abstract thought evolved within the context of ancestral abilities for interacting with the world, namely selection among competing action plans. Therefore, studying perceptual judgments in mice can uncover the neural mechanisms that preceded the emergence of abstract thought.

## Materials and Methods

### Animal subjects

All surgical and behavioral procedures were approved by the Institutional Animal Care and Use Committee. Experiments were conducted with male and female mice between 8 and 24 weeks of age. Four transgenic strains purchased from the Jackson Laboratory were used to create the mouse lines used for optogenetics and calcium imaging: PV-Cre (JAX #008069), RCL-ChR2 (also known as Ai32; JAX #012569), CaMKII-tTA (JAX #007004), and TRE-GCaMP6s (JAX #024742). All mouse strains were of C57BL/6J background. For optogenetics we used 10 PV-ChR2 mice in behavioral experiments and an additional 4 PV-ChR2 mice for electrophysiological measurements, for widefield imaging we used 11 CaMKII-GCaMP6s mice, and for two-photon imaging we used 13 CaMKII-GCaMP6s mice. No statistical methods were used to predetermine the sample size. All mice undergoing training were housed alone under an inverted 12-hour light/dark regimen and trained in darkness during the dark part of their cycle.

### Surgical procedures

All surgeries were performed under 1–2% isoflurane anesthesia. After the mouse was anesthetized, its scalp was shaved and lidocaine ointment (Emla) was applied to its skin. A circumferential incision was then made to expose as much of the skull as possible. For optogenetics and widefield imaging, after clearing the skull’s periosteum, two small marks were made using nail polish at the positions of bregma and lambda for mapping purposes. We then applied a layer of cyanoacrylate (Krazy Glue) to optically clear the bone and placed a 3-mm-diameter coverslip (thickness = 150 μm) to make the surface even. This resulted in clearly visible blood vessels after a few minutes. A headbar was then affixed behind the coverslip using more cyanoacrylate. Finally, a black, 2-mm-high light shield was glued in front of the coverslip to attenuate the light from the optogenetics or widefield imaging reaching the mouse’s eyes.

For two-photon imaging, after clearing the skull’s periosteum, we performed a 3 mm craniotomy using a cranial punch, centered at either −1.5 AP, 3 ML (for S1-C2 imaging), −1.5 AP, 1.75 ML (for RSC and PPC-A imaging), or 1.9 AP, 1.15 ML (for MM and ALM imaging). We then constructed an imaging window from either one (for S1-C2, RSC, and PPC-A) or three (for MM and ALM imaging) layers of 3 mm diameter coverslips (Harvard Apparatus). Animals were monitored for one week of recovery before training began.

### Behavioral apparatus, training, and stimulus delivery

The behavioral setup was based on a custom LabVIEW program controlling and receiving data from both a DAQ card (PCIe 6351, National Instruments) and three Arduinos (Mega 2560). The three Arduinos were used to create a pulse-width modulated output for the servomotor and speaker, to control the linear motors, and to synchronize the LED colors with the frames during widefield imaging (see “Widefield imaging”). 25 mice were trained in the blocks task, and 9 mice were trained on the lick-left/lick-right detection task with a delay. Trials were automatically initiated with a 1–2 second baseline period (randomized in each trial) prior to the tone. The tone was a 100-ms beep coming from a small speaker behind the mouse (equidistant from both sides) at 5 kHz and ∼70 dB at the mouse’s position. In go trials in the standard blocks, and no-go trials in the reversed blocks, the tone was concurrent with a galvanometric stimulation of the C2 whisker. This stimulation was done by inserting the C2 whisker into a 23-gauge blunted needle attached to the galvanometer. To ensure stimulation reproducibility, we positioned the tube tip 2 mm from the whisker’s base using an XYZ-adjustable stage. Finally, we reversibly attached the whisker to the tube using uncured silicone (Yellow Body Double, Smooth-On).

The lick ports consisted of two 18-gauge blunted needles mounted on rotary servomotors (Turnigy TGY-D003iV), which themselves were mounted on linear actuators (DC House L11010101011-1). At the end of the tone, the lick ports rotated toward the mouse^35^. This was done to prevent any confounds from early licking before or during the stimulation. The servos-lick ports complex was mounted on an XYZ-adjustable stage, and positioned for each mouse such that at the end of the servo’s movement, the lick ports were 2 mm below the upper lip, 1 mm posterior to the tip of the lower lip, and approximately 1 mm towards either the left or right side. The mouse had a 500-ms response window to lick before the servo rotated the lick ports away. Either a piezo or capacitive sensor was attached at the base of the lick ports and sensed the licks, transmitting them to the DAQ card. If the mouse licked, then depending on whether it licked the correct or incorrect lick port, it respectively received a reward or a punishment upon the end of the response window. The reward consisted of a 7 μL drop of 5% sucrose water, while the punishment was a 500-ms airpuff to the face. If the mouse was rewarded (hit), then the lick port stayed for an additional 4–6 seconds to allow the mouse to fully consume the water. If it was punished (false alarm), then there was an additional 5–8 second timeout period. Otherwise (miss or correct rejection), the delay to the beginning of the next trial was 2 seconds. In each case, we also added a random 1–2 second variable delay from trial start until tone onset to prevent mice from predicting it.

Animals were trained for approximately one month in either the blocks or the delayed lick-left/lick-right paradigm. One week prior to the beginning of training, they were water-restricted and habituated to the setup by being provided free water on the setup. Eventually, a trial structure was introduced such that all trials were go trials with a single lick port and the mice were taught to lick after the tone. Once the mice were habituated, all whiskers on the left and right whisker pads were trimmed other than C2 on both pads. After mice learned to lick a single lick port, two lick ports, one on the left and one on the right, were introduced to familiarize mice with the mechanics of licking right vs. left. Initially, only the lick port corresponding to the correct choice in the lick-left/lick-right task was presented on each trial (i.e., all trials were go trials), and when mice licked that lick port then the other lick port was presented on the next trial. Once we observed mice learned to lick to either side, both lick ports were introduced (day 1 on the learning curves in Extended Data Fig. 1b). During 14 days of training, mice were trained to perceive a relatively strong right C2 whisker stimulation (∼6° amplitude, 20 Hz, 100 ms). In the blocks paradigm, stimulus trials were lick left trials and no-stimulus trials were lick right trials. In the delayed lick-left/lick-right paradigm, the same contingency held for the congruent-trained mice, and the opposite contingency (stimulus → lick right, no stimulus → lick left) held for the incongruent-trained mice. Initially, lick right/left trials were not random, but required the mouse to perform correctly either once or twice (randomized to prevent pattern learning) before switching. Only once the mice reached *d’* = 1.5 did we make lick right/left trials random, at which point we observed that doing so had no detrimental effect on performance. After 7 days of training for each mouse, we proceeded to the next stage. For the lick-left/lick-right mice with a delay, this was the introduction of a delay. For the contingency blocks mice, this was the introduction of the block structure, wherein for 50 trial blocks one lick port was retracted by 5 mm using the linear motor, just out of the reach of the mouse’s tongue. To minimize any differences in sounds across blocks, both lick ports were still rotated using the servomotors toward the mouse. After an additional seven days of training for each mouse, we performed psychometric testing (four sessions in the blocks task, and two sessions in the delayed lick-left/lick-right task) to determine the perceptual threshold (see “Psychometrics” section). Finally, we either recorded from the mouse (see “Widefield imaging” and “Two-photon imaging” sections) in the task or performed optogenetic inhibition (see “Optogenetics” section). In recording sessions, in 50% of trials the mouse was given no stimulus, in 40% of trials it was given the threshold stimulus, and in 10% of trials it was given the strong stimulus. The reason for this ratio is that we largely wanted to concentrate on threshold stimulus trials as they provided us with similar numbers of stimulus present and stimulus absent judgments, while strong stimulus trials were largely used for control purposes. In inhibition sessions, either the same procedure was followed (cortex-wide inhibition) or in 50% of trials the mouse was given no stimulus and in 50% of trials it was randomly given one of six psychometric stimulus intensities (targeted inhibition). Most (seven of nine) mice trained on the delayed lick-left/lick-right task then proceeded to learn the opposite contingency.

### Psychometric measurements and simulations

During learning, the discrimination index (*d’)* was calculated for each session as *Z*(*P*(hit)) − Z(*P*(false alarm)), where Z(*P*) is the inverse of the standard normal cumulative distribution function (cdf), evaluated at the probability *P*. Intuitively, it represents the normalized difference between the hit percentage and false alarm percentage, expressed in standard deviations.

After learning each task but before recording, the stimulus amplitude was varied to obtain lick percentages across intensities. To identify the best fitting psychometric function, we compared the full logistic and Weibull fits from the Python library psignifit to partial logistic fits using least-squares regression (Extended Data Fig. 1f) by calculating both the likelihood and Bayesian information criterion (BIC) of each fit across both standard and reversed blocks. Based on the findings, we ultimately used partial logistic regression with two separate slopes and biases for the contingency blocks and lick-left/lick-right without delay tasks, and full logistic regression with added guess and lapse terms for the lick-left/lick-right with delay task. The intensity threshold was obtained from these fits, and this was the main intensity given to the mice in the subsequent cortex-wide inhibition or recording sessions. For chronometric fitting (Fig. 1d), we applied a simple linear regression.

To model how the psychometric curves are affected by changes in bias and sensitivity, we began by performing logistic regression on the control psychometric data of the optogenetic mice to obtain the empirical biases and sensitivities for each block (Extended Data Fig. 3a). We then applied shifts to either the empirical sensitivities or biases, across both blocks, in steps of 0.05 from −1 to 1. From the resulting set of shifted parameters, we then used the logistic function to obtain theoretical psychometric curves, consisting of lick rates for seven stimulus intensities between 0 and 1 for each block. We subsequently performed 1000 simulations of 1000 trials each with these probabilities, such that half the trials were standard and half were reversed, with half of each being no-stimulus trials and half distributed among the remaining six intensities. The same simulations were also applied to a no-shift control. Then, we fit logistic curves on the simulated trial data, and calculated the following metrics relative to the control curves: (1) standard false alarm (FA; lick rate at zero intensity) shift, (2) standard sensitivity shift, (3) both blocks (averaged) sensitivity shift, (4) horizontal shift in intersection, (5) standard bias shift, (6) both blocks (averaged) bias shift, and (7) vertical shift in intersection. In each case, we calculated the discrimination index (*d’*) between the distributions of each metric under sensitivity vs. bias shifts (Extended Data Fig. 3c) and each kind of shift vs. control (Extended Data Fig. 3d,e).

### Video monitoring

Two CMOS cameras (DMK 37BUX287) were used to monitor mouse movements. They were positioned to capture the right side of the mouse’s face and its body. Footage was either recorded at 300 frames per second (fps) and down-sampled to 30 fps, or recorded directly at 30 fps. To illuminate the mouse, we placed one 850 nm LED (M850L3, Thorlabs) in front of the mouse and another 850 nm LED underneath it. Other sources of background illumination (especially the 470 nm and 530 nm light during widefield imaging) were removed by placing 830–865–nm band-pass filters (BP845, MidOpt) in front of the sensor in each camera.

### Optogenetics and extracellular recording control

Optogenetic inhibition was performed by activating ChR2-expressing inhibitory (PV) neurons in transgenic mice using a 447 nm laser (CNI) coupled to a galvanometric scanning photostimulation system^18^. While near-ChR2 excitatory peak wavelengths (e.g., 473 nm) are more typically used, the excitation spectrum of ChR2 is broad enough that 447 nm is at ∼0.95 of the peak^90^. Importantly, however, neural tissue penetrance at this wavelength is about 50% of that at 473 nm^91^. We coupled the laser to a 200 μm multimode fiber which was directed via an achromatic collimator (F950FC-A, Thorlabs) to a scanning galvanometric mirror system (GVS002). From there, the beam was focused obliquely onto the surface of the brain with an LSM03-VIS scan lens (Thorlabs). As a result of multimode fiber limits, the resulting beam diameter was considerably larger (estimated at ∼1 mm FWHM using the widefield camera) than when using direct excitation (measured at 0.27 mm FWHM in ref. 92).

Laser intensity and position were controlled with custom Python software and a data acquisition card (PCI-6713, National Instruments). The laser was pulsed at 40 Hz as a square wave with an 80% duty cycle, and set to 5 mW overall from 200 ms before tone onset until the end of the response window, when it linearly tapered off to 0 mW over 200 ms to minimize rebound effects. When stimulating more than one cortical region, the laser position moved between the regions at 200 Hz (overlaying the 40 Hz square pulses) with the laser turned off during the movement. In that case, due to power loss as each region was stimulated for shorter durations, laser power was increased to maintain 5 mW on each region overall.

To target the laser to the intended regions on the skull, we used the widefield camera described in “Widefield imaging”. After focusing the laser on the cortex and verifying construct expression (via the ChR2-coupled YFP in these transgenic mice), we rigidly aligned the cortex to the Allen CCF v3 as also described in “Widefield imaging”. In addition, we targeted a region outside the cortex for control stimulation. In the cortex-wide inhibition experiments (Fig. 2c,e, Extended Data Fig. 2e), we inhibited one of 42 locations across the two hemispheres in 80% of the trials and the control region in the remaining 20%. These locations were separated by 1.15 mm horizontally and 1.4 mm vertically, with an additional vertical shift of 0.25 mm in lateral regions to include the center of S1-bc among the targets. In the targeted inhibition experiments (Fig. 2f), we inhibited one of eight targeted regions in 40% of the trials and the control region in the remaining 60%. The percentage of control trials was lower in the targeted inhibition compared to cortex-wide inhibition, as our manipulation more often affected behavior and we sought to avoid confusing the mouse.

In the extracellular recording control experiments, we manually marked points adjacent to an axis spanning from the center of S1-bc to the targeted inhibition point in M2, from 0 to 3 mm at 0.5 mm intervals. We then inserted a four-shank Neuropixels 2.0 probe with a channel configuration that spanned either 200 to 700 μm (two mice) or 300 to 800 μm (two mice), and delivered laser pulses with the same parameters as described above for 250 trials, randomizing the inhibition distance on each trial. The data was band-pass filtered between 300 and 6000 Hz and median-subtracted. It was then spike sorted using Kilosort 4^92^ and automatically curated with UnitRefine^93^, followed by manual curation to remove any artifacts caused by the laser stimulation. Putative inhibitory and excitatory units were defined as those with peak-to-valley times below and above 0.475 ms (selected based on the separation of two peaks in the bimodal distribution of peak-to-valley times), respectively. Finally, we averaged the activity of each cell in a window of 800 ms before each inhibition, and the window of 800 ms during the inhibition, and obtained the average % decrease across the excitatory population and % increase across the inhibitory population.

### Widefield imaging

Widefield imaging was done using a macroscopic lens (Nikon AF Micro-NIKKOR 60 mm f/2.8D) mounted on a scientific complementary metal-oxide semiconductor (sCMOS) camera (Quantalux, Thorlabs) running at 60 frames per second. To capture GCaMP fluorescence, a 500 nm long-pass filter (FELH0500, Thorlabs) was placed in front of the camera’s sensor. The total field of view was 11.5 x 9.5 mm and the image resolution was 230 x 190 pixels after 4x spatial binning (spatial resolution: ∼50 μm per pixel). A “dark frame” was collected at the beginning of each session from 100 frames in the absence of light and subsequently subtracted from all image frames. To isolate Ca^2+^-dependent signals, we corrected for intrinsic signals by multiplexing excitation light at two different wavelengths: 470 nm (blue, M470L3, Thorlabs) and 530 nm (green, M530L3, Thorlabs). This was accomplished by coupling the two LEDs into the excitation path using a 505 nm long-pass dichroic mirror (DMLP505R, Thorlabs), and using the strobe counter output of the camera to alternate illumination between the two LEDs from frame to frame. This resulted in one set of frames with blue and the other with green light at 30 frames per second each. Now, whereas the signal from the blue light depends multiplicatively on both Ca^2+^ activity and intrinsic signals, specifically blood volume, the signal from the green light depends only on intrinsic signals^94^. Therefore, we could approximately remove the intrinsic signals from each blue frame by dividing by the interpolated green frame:

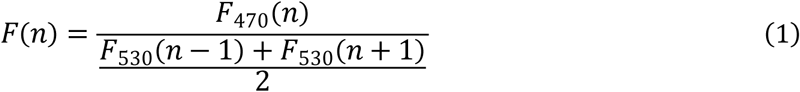

where *n* is the frame number^94^. This ratio was then normalized to its mean during the baseline period of each trial, defined as the 200 ms immediately preceding tone onset. Next, all frames were motion-corrected in the *x* and *y* axes using FFT-based subpixel registration^95^. Finally, they were rigidly aligned to the Allen CCF v3^96^ using the locations of lambda and bregma, marked on the skull during the surgery stage, and the locations of the left and right hemisphere C2 barrels, found by stimulating the left and right C2 whiskers following the first recording session. Alignment was tested by extending the tone durations to one second and observing activation of the left and right primary auditory cortices in their expected location on the Allen CCF.

### Two-photon imaging

Two-photon calcium imaging was performed with a commercial two-photon microscopy system (Bruker Ultima). We recorded from one of five regions in the hemisphere contralateral to the stimulus: RSC (retrosplenial cortex), PPC-A (posterior parietal cortex, anterior area), MM (medial motor cortex), ALM (anterolateral motor cortex), and S1-C2 (C2 barrel of the somatosensory cortex). The first four were targeted by stereotaxic coordinates (RSC: −1.4 AP, 0.9 ML; PPC-A: - 1.65 AP, 1.8 ML; MM: 1.3 AP, 0.8 ML; ALM: 2.5 AP, 1.5 ML; distances in mm) while the last was targeted by performing widefield calcium imaging over a −1.5 AP, 3 ML craniotomy and finding the peak of the C2 whisker response. We imaged either S1-C2, or RSC and PPC-A, or MM and ALM in separate groups of mice. From each region, we recorded at 30 Hz from a 500 × 500 μm area using an 8 kHz resonant laser scanner. Excitation was via a Ti–sapphire laser (Newport) tuned to 920 nm with a 20× (1.0 NA, Olympus) water-immersion objective. Power out of the objective was controlled by calibrated rotations of a half-wave attenuator and depended on the magnification of the scan but was typically 20–40 mW. The excitation signals from GCaMP were passed through a dichroic mirror (HQ575dcxr, Chroma) and amplified by a GaAsP photomultiplier tube (Hamamatsu).

We recorded from layers 2–4 for each mouse and region, at depths ranging from 160 to 470 μm from the dural blood vessels. For each recorded region, we performed preprocessing in suite2p^97^, which included further (local) motion correction, ROI detection, and subtraction of surrounding neuropil signals. Neurons were curated manually from the suite2p results. Prior to all subsequent analysis steps, the fluorescence traces were normalized to the *F0* value in each trial, taken to be the mean of 500 ms prior to the tone onset, by calculating *(F(t)-F0)/F0*. For classification analyses of block and direction coding, we also used the deconvolved spike estimates from suite2p.

### Selectivity index analysis

We used selectivity index (SI) analysis across matched trial types to compute the magnitude and significance of coding of different variables. This approach is a generalization of classical choice analysis^98^ except combining multiple trial-type comparisons. We will describe this method for each type of activity we analyzed in this work (pixels, regions, and cells).

The procedure of this analysis can be seen as constructing two distributions of single-trial activities, and then calculating a measure of the distance between them. In particular, we calculated the Mann–Whitney *U* across activities from two trial types with *n*_1_ and *n*_2_ trials each, then computed the SI using the formula below:

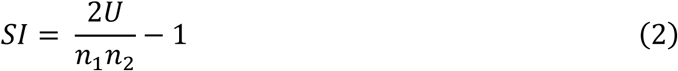

SI has several useful properties: (1) it is 0 for perfectly overlapping distributions, (2) it ranges between −1 and 1, and (3) it is independent of the number of trials of each trial-type, allowing for imbalances. For widefield imaging analysis, the distributions were either separately constructed from the activities of each widefield pixel averaged over each time bin (i.e., 0 to 100 ms, 100 to 200 ms, etc.; Fig. 3c,f,g) or the mean activity of all pixels in each region at each frame (i.e., 0 ms, 33 ms, etc.; Fig. 3e). For two-photon imaging analysis (Fig. 4g,h), the distributions were separately constructed from individual cell activities averaged over the entire tone/stimulus and response periods.

For calculating the stimulus response in the standard block (Fig. 3c), we compared standard misses (stimulus given, no lick, stimulus absent judgment) to standard CRs (no stimulus given, no lick, stimulus absent judgment) and standard hits (stimulus given, lick left, stimulus present judgment) to standard FAs (no stimulus given, lick left, stimulus present judgment), while in the reversed block, we compared reversed CRs (stimulus given, no lick, stimulus present judgment) to reversed misses (no stimulus given, no lick, stimulus present judgment) and reversed FAs (stimulus given, lick right, stimulus absent judgment) to reversed hits (no stimulus given, lick right, stimulus absent judgment). Hence, in each case we compared the distribution when the stimulus was present to that when it was absent, controlling for the action in each comparison (either both no lick, or both lick). For each comparison, we created separate distributions for the activity of each pixel at each time bin. We then calculated the overall SI using a weighted average across comparisons (specifically, weighted by the product of the number of trials, *n1n2*). For the overall stimulus response calculation (Figs. 3e, 4g), we calculated the weighted average across all four comparisons.

The SI computation for block and direction was similar, but was based on different trial types. For block, we compared standard misses (stimulus given, no lick, stimulus absent judgment) to reversed CRs (stimulus given, no lick, stimulus present judgment) and standard CRs (no stimulus given, no lick, stimulus absent judgment) to reversed misses (no stimulus given, no lick, stimulus present judgment). For direction, we compared standard hits (stimulus given, lick left, stimulus present judgment) to reversed FAs (stimulus given, lick right, stimulus absent judgment) and standard FAs (no stimulus given, lick left, stimulus present judgment) to reversed hits (no stimulus given, lick right, stimulus absent judgment). Note that while the matching procedure in this case was controlled for stimulus and action, it could not control for perceptual judgment coding, which we found to be negligible (Fig. 4). In addition, any block coding was subsumed in the direction coding. Thus, we excluded all cells found to be significantly block-coding using the below procedure from the direction coding subset prior to analysis (Extended Data Fig. 9f).

Significance in each case was calculated by comparing the measured SIs to the distribution of SIs calculated from randomly shuffling trial labels 1000 times. Specifically, for each pixel or cell, we randomly set trials as belonging to either the first or second trial type being compared regardless of their actual trial type to simulate what we would expect by chance. Finally, we applied the Benjamini–Hochberg procedure for false discovery rate (FDR) correction of the resulting *P* values at α = 0.05.

### Linear (ridge) regression model

We sought to create a model of either widefield calcium activity (Fig. 3i,j) or single-cell calcium activity (Fig. 4i) given by *Y* = *X β* where *Y* is either the widefield activity matrix or single-cell activity matrix, *X* is a design matrix consisting of task and behavioral factors, and *β* is the kernel matrix that relates *X* to *Y*. Our goal was to evaluate how much each task and behavioral factor uniquely contributes to the model’s predictive performance of activity in each pixel or neuron.

To create the widefield activity matrix *Y*, we first concatenated the *(F(t)-F0)/F0* data containing the pre-stimulus baseline up to the end of the response window (to exclude confounding contributions from rewards and punishments) for each mouse across all its recorded sessions, including both blocks. We then reduced the resulting matrix for each mouse to its 113 highest-varying components (selected as the number of components that explains 99% of the variability in the data on average across mice) using singular value decomposition (SVD). SVD returned both spatial components *U* (sized atlas height by atlas width by number of components), temporal components *V*^*T*^(sized number of components by number of frames), and singular values *S* (sized number of components by number of components). To prevent overfitting and reduce computational complexity, the subsequent regression was performed on the transpose of the product *SV*^*T*^ as our *Y* (sized number of frames by number of components). As a sanity check, we verified we could approximately reproduce the original widefield *(F(t)-F0)/F0* videos by convolving these components with the spatial ones (data not shown). To create the single-cell activity matrix *Y*, we concatenated the *(F(t)-F0)/F0* values of each recorded cell for each session such that *Y* was sized number of frames (concatenated from all trials, pre-tone until the end of the response window) by number of cells.

To create the design matrix *X* (sized number of frames by number of variables) for both widefield and single-cell models, we first chose predictive task (whisker stimulus, action, direction, perceptual judgment, tone, block identity) and behavioral factors (face and body motion energies). These factors were either analog (face and body motion energies), ternary (direction, i.e., 1 for lick left, −1 for lick right, and 0 for no lick), or binary (the remaining ones). To capture delayed and biphasic responses, we also created time-shifted copies of the binary event factors. For example, for the whisker stimulus, we created a vector consisting of 0s and 1s across all frames (concatenated across all trials) recorded for each mouse. Vector elements corresponding to frames during which the stimulus was given were set to 1 and all other elements were set to 0, and this vector was the first column in the design matrix. Then, we shifted this vector forward by one index (a frame, equivalent to 33 milliseconds) and placed this new vector as the second column in the design matrix. We continued adding shifted copies of the stimulus vector to the design matrix until the shifted vectors spanned the entire duration of the response window (0–600 ms from stimulus onset). A similar procedure was performed for the tone and perceptual judgment (according to the table in Fig. 1c), in each case adding more columns to the design matrix consisting of each factor’s original vector along with its shifted copies, with the latter two factors shifted to span until the end of each trial. We did not shift the analog movement vectors due to their already highly auto-correlated nature, and the resulting diminishing returns on explained variance given high computational demand (data not shown).

For the action and direction variables, we first added a column to the design matrix with elements corresponding to lick events being set to 1 for action, and either 1 (lick left) or −1 (lick right) for direction, and all other elements set to 0. We then added shifted vectors to span from the lick event until the end of the response window. In addition, we shifted the lick vector back by up to 500 ms to account for motor preparatory neural activity and added these backward-shifted copies to the design matrix as well. The block identity variable was set to 1 for all frames in the standard blocks and −1 for all frames in the reversed blocks.

For the uninstructed face and body movement variables, we first concatenated the frames from the face and body cameras across all trials and then calculated the absolute differences between the pixels of successive frames (i.e., motion energies). We then performed a similar procedure to that for widefield activity above. Namely, we reduced the resulting matrices to their 167 and 186 highest-varying components (again selected as the number of components that explain 99% of the variance in the data) using singular value decomposition (SVD) and included their temporal component product *SV*^*T*^ as columns in the design matrix. We again verified we could accurately reproduce the original movement videos from these components (data not shown).

Importantly, because some task variables like action and direction could be inferred from video data, we needed to ensure their explanatory power did not overlap with other model variables^35^. To do so, we performed QR decomposition of a reduced design matrix *X_r_* containing instructed movement variables as well as the face and body movement variables. The QR decomposition spanned the same space as *X_r_*, but such that all the columns (i.e., the variables) were orthogonal to one another. Finally, we replaced the uninstructed movement columns of the full design matrix *X* with the corresponding columns of the QR-decomposed reduced design matrix. This allowed the model to improve the fit to the data using any unique contributions of the uninstructed movement variables while ensuring that the weights given to other variables were not altered.

To obtain *β*, the kernel matrix that relates *X* to *Y* (sized number of components by number of frames), we used ridge regression^35^. A ridge penalty was optimized for all components of widefield data or all cells in the single-cell data using SciPy’s fminbound function over the [0, 30000] range with tenfold cross-validation. We then applied this ridge penalty when fitting the kernels for the remaining data via fivefold cross-validation. At each fold, we set aside test data and calculated *β* using the formula for linear regression with ridge regularization parameter λ:

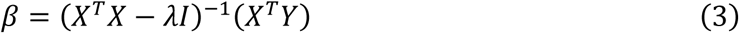

where I is the identity matrix. We then calculated the Pearson correlation coefficient between the modeled *Y* (*X β*) of the test data and its real value, squared it, and averaged across the folds to obtain the cross-validated explained variance (cv*R*^2^).

To assess the unique explained variance of each individual factor (Δcv*R*^2^), we removed this factor as well as all its shifted copies from the design matrix and re-fit the data, giving us the explained variance of the reduced model for that factor. We then subtracted the explained variance of the reduced model from the explained variance of the full model to obtain Δcv*R*^2^, the unique explained variance of that factor. Importantly, this approach allowed us to assess the contribution of variables in a more specific way than SI analysis. For instance, given that licking was associated with more uninstructed movements (Fig. 1f), the difference between lick and no-lick trials gave us the neural correlates of both. In contrast, in the regression model we could obtain the unique explained variance of licks independently of the uninstructed movements (which were orthogonalized with respect to licks) and vice versa.

### Population classification

To ask how the different variables are encoded by simultaneously recorded single-cell populations of individual areas (Fig. 4j and Extended Data Fig. 8), we used a cross-validated classification approach. First, we needed to decide what trials fall into each category. Because simply taking all trials with or without a certain variable (e.g., stimulus) could result in imbalances in other variables (e.g., action) that would confound the result, we used a similar approach to the trial type matching described above for SI analysis. However, as in this case we could not do a weighted mean of classifier performance across each comparison (as the classifier could be different in each case, while we want it to generalize), we applied a different approach: we calculated which trial type had the minimum number of trials, then randomly subsampled (under 10 different initializations) this number of trials from all other trial types. This ensured that all other variables than the one being classified were balanced, and allowed us to apply a general classifier to all trial types based on the presence or absence of the variable.

For stimulus (separately across threshold and strong stimulus trials), block, and direction classifiers, the trial types used are described in the “Selectivity index analysis” section above. For the action classifier, we compared [standard hits, standard FAs, reversed FAs, reversed hits] to [reversed CRs, reversed misses, standard CRs, standard misses], which, combined with the above subsampling procedure, controlled for any stimulus-, direction-, judgment-, and block-related differences. For the action-independent judgment classifier, we compared [standard hits, reversed CRs, standard FAs, reversed misses] to [reversed FAs, standard misses, reversed hits, standard CRs], which similarly controlled for any differences other than in judgment.

We created a matrix sized trials by cells for each session and tone-aligned frame (e.g., 0 ms, 33 ms, etc.) which we filled with the *(F(t)-F0)/F0* values from trials of each subsampling initialization. We then also created a target vector consisting of 0s or 1s indicating the absence or presence of the variable for each subsampled trial (e.g., 0 for selected trials without a stimulus, and 1 for selected trials with a stimulus). Subsequently, we trained various decoders on columns of the feature matrix and target vector, namely either logistic classification with L1 regularization (Fig. 4j) or other classifiers (labeled in Extended Data Fig. 8). In each case, we used stratified fivefold cross-validation to compute decoder performance (AUC), wherein trials were randomly split into train and test sets consisting of 80% and 20% of the data, respectively, such that the variable identities (i.e., 0s and 1s) were balanced in each split. The L1, L2, and elastic net (half L1, half L2) penalties, if used, were chosen from the set [0.001, 0.01, 0.1, 1, 10] using a separate cross-validation to find the penalty that yielded the highest AUC across the test sets. Finally, we averaged AUCs across all 10 random subsampling initializations.

To test the generality of population classifiers across trial types (Fig. 4k), we formed two balanced sets of train and test trials for the judgment and strong stimulus variables, with the latter used as a control. For judgment, we trained an L1-regularized logistic classifier to classify the set [standard hits, reversed misses] (stimulus present judgments) and [standard CRs, reversed FAs] (stimulus absent judgments), which was balanced in stimulus, block, and action via the above subsampling procedure, and tested it on classifying the set [reversed CRs, standard FAs] (stimulus present judgments) and [standard misses, reversed hits] (stimulus absent judgments), which was similarly balanced. We then switched the training and testing sets, and averaged the resulting AUC at different times with respect to stimulus onset. For strong stimulus, we similarly trained a classifier to classify the set [standard hits, standard misses], restricted to strong stimulus trials, and [standard FAs, standard CRs], and tested it on classifying the set [reversed FAs, reversed CRs], again restricted to strong stimulus trials, and [reversed hits, reversed misses]. We then switched the training and testing sets as for judgment.

Lastly, we calculated the parallelism score (PS) across the same sets for judgment and strong stimulus. To do so, we first trained an L2-regularized logistic classifier (L1 regularization was not used here due to its tendency to sparsify weights, which distorts angle measurements) on each of the aforementioned training/test sets independently. We then computed the parallelism score as the cosine angles across the two sets of weights:

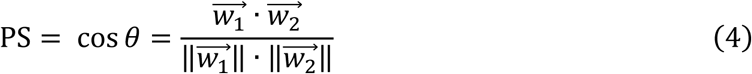

where 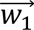 is the weights vector of the classifier trained on the first train/test set from before and 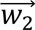 is the weights vector of the classifier trained on the second.

### Network model

We created three drift-diffusion neural network models (Fig. 6a, Extended Data Fig. 10a,c). In the first (‘action-coupled’) model, a population of 100 stimulus coding units and 100 block coding units synapsed onto two populations of direction units, namely 100 lick left and 100 lick right units. Each unit was a simple scalar with activity between 0 and 1 in each millisecond. The stimulus coding units had a synaptic strength determined by the sensitivity in the current block (*S_s_* in the standard blocks and *S_r_* in the reversed blocks), while the block coding units had unitary synaptic strength. In the first 100 ms, the sum of stimulus unit activity during each millisecond was given by *s*, while the sum of block unit activity was *b_s_* in the standard blocks and *b_r_* in the reversed blocks. Additionally, we added a Gaussian noise input with zero mean and standard deviation *σ*. In the following 500 ms (i.e., the response window), only the noise component remained. This activity (i.e., the sum of stimulus, block, and noise inputs) was integrated over time to determine the lick left and right unit activity output. Lick left and right units followed a competitive principle: positive inputs excited lick left units but inhibited lick right units, while negative inputs inhibited lick left units and excited lick right units. If the summed activity of either lick left or lick right units reached the bound given by *B,* a lick in the respective direction was triggered after NDT (non-decision time) milliseconds. If a lick did not occur within the response window of 500 ms, the trial was registered as a no-lick trial. In the second (‘action-coupled, block-unit free’) model, we replaced the block units with bias units with activity *b* in both blocks, but otherwise the model behaved identically. In the third (‘action-independent’) model, we kept the replacement of block units with bias units as in the second model, and made three additional alterations: the two populations of direction units were replaced by a single population of 100 judgment units (also with a threshold *B*, but no competition with another population), an additional free hazard time parameter *Th* was added that allowed for earlier response if this threshold was not reached (to allow mice to report ‘stimulus absent’ judgments before the end of the response window), and a population of block units downstream of the judgment was included that allowed for the judgment to be translated into different actions depending on the block.

To fit the models to the psychometric and chronometric curves, we optimized their parameters (in the first model: *S_s_*, *S_r_*, *b_s_*, *b_r_*, *σ*, *B*, and NDT; in the second: *S_s_*, *S_r_*, *b*, *σ*, *B*, and NDT; in the third: *S_s_*, *S_r_*, *b*, *σ*, *B*, NDT, and *Th*). For this purpose, we were able to fully ignore the individual units and focused on the accumulation process (i.e., only the total inputs from each unit population were considered). In the first and second models, due to the competitive principle explained above, we were able to model the difference between the lick left and lick right populations (i.e., sum of lick left unit activity – sum of lick right unit activity) as a drift-diffusion process with two bounds and a step-function drift rate. As no analytic solution of such a process was available, we used the open-source Python library PyDDM^99^, which could solve it numerically using the Fokker-Planck equation. In particular, we set up 14 conditions (seven stimuli in each of two blocks), and minimized the squared error loss across both the psychometric and chronometric curves relative to the real data using differential evolution.

In the third model, as there was only a single (upper) bound, we were able to derive an analytic solution using known equations governing Wiener processes (Brownian motion) with an absorbing bound. Specifically, a process with constant drift rate given by *v* = *s*\**S* + *b* (where *s* is the stimulus intensity, *S* is sensitivity, and *b* is bias), noise *σ*, and bound *B*, will have a first-passage density (probability density of being absorbed at the upper boundary) given by (equation 1 in ^100^):

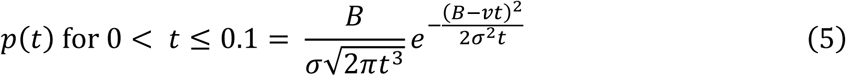

Let *T* denote the first-passage time to the upper boundary. We can then obtain the total probability *P*(*T*≤0.1) of this first-passage time within the first 0.1 seconds by numerically integrating equation (5) between 0 and 0.1 seconds, and the expected absorption time by computing:

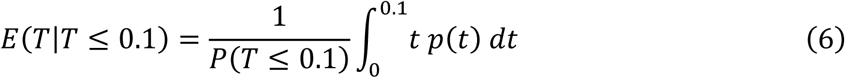

Next, we turn to the probability and expected absorption time in the diffusion-only (*v* = 0) phase between 0.1 and 0.6 seconds after stimulus onset, assuming an absorption did not occur. Given a boundary at *B*, the probability density of a Wiener process at time *t* with respect to the integrated evidence *y* is given by (modified from the transition density in Appendix 1, section 17 in ^101^, with speed measure multiplier *σ* instead of 2):

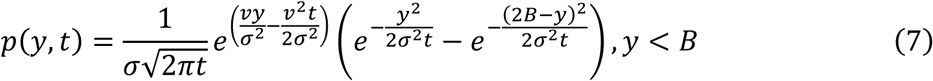

Given starting point *y*(*t* = 0.1) *= Y_0_*, the first-passage density of a diffusion-only Wiener process is given by our equation (5) with *v* = 0, *B* changed to *B* – *Y_0_*, and t shifted by 0.1 seconds:

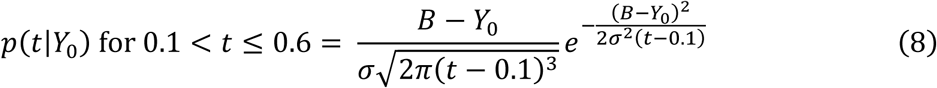

Combining the two, the conditional probability density of first passage between 0.1 and 0.6 is given by integrating the product of the probability density in equation (7) over all positions at 100 ms and the first-passage density of each position in equation (8):

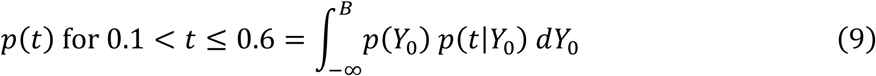

where *p*(*Y*_0_) = *p*(*y*, *t* = 0.1). The total probability *P*(0.1 < *T* ≤ 0.6) of absorption during the lick window of 0.1 to 0.6 seconds is then given by integrating this density between *t* = 0.1 and 0.6, and the expected absorption time is given by the integral:

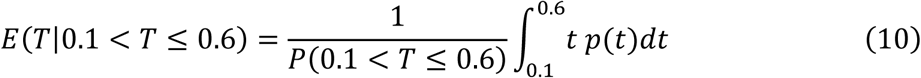

Overall, combining both time periods, we obtain

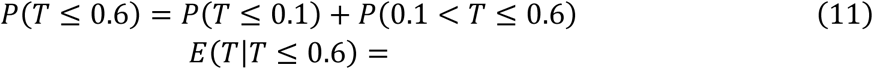

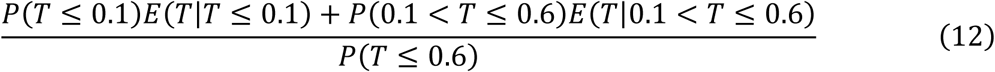

Finally, we accounted for the non-decision time (NDT) by adding it to the expected lick time, and replaced the limit of the response window (0.6) with the aforementioned free hazard time parameter *Th*. The resulting absorption probability *P* and mean absorption time *E*(*T*) were computed separately across blocks and stimulus intensities and their squared error loss relative to the real psychometric and chronometric curves was minimized using SciPy’s basin-hopping algorithm followed by L-BFGS-B gradient descent.

To simulate neural data in the first model (Fig. 6e,f), we ran Monte Carlo simulations of the full network model with the mean of the obtained macroscopic parameters (Fig. 6b) for a single, 350-trial session (a typical number of trials in the real sessions used to obtain the data in Fig. 4g,h). As an additional source of variability, each unit was chosen to have a random level of responsiveness on a trial-by-trial basis, ranging uniformly from 0% to 100%. The activity of each unit was derived from dividing its respective macro parameter by the number of active units on any given trial (e.g., if 50% of the 100 stimulus units were active and the stimulus input was 0.5, then each active stimulus unit had magnitude 0.5/(0.5*100) = 0.01). We then performed exactly the same SI analysis on all the model units as for the processed two-photon data.

### Analysis software

All post-preprocessing analysis and visualization of behavioral, optogenetic, widefield, two-photon, and model data were performed using custom scripts in Python 3.8 or 3.10, as well as the following third-party libraries: ipywidgets, NumPy, Matplotlib, Pandas, scikit-learn, SciPy, statsmodels, opencv-python, PyDDM, and psignifit.

## Data and code availability

Processed data and code will be publicly deposited upon publication and are available upon request in the meantime.

## Acknowledgments

We thank S. Musall, D. Rinberg, O. Barak, N. J. Priebe, and M. Diamond for valuable discussions and suggestions about this work.

## Extended Data

**Extended Data Figure 1:**
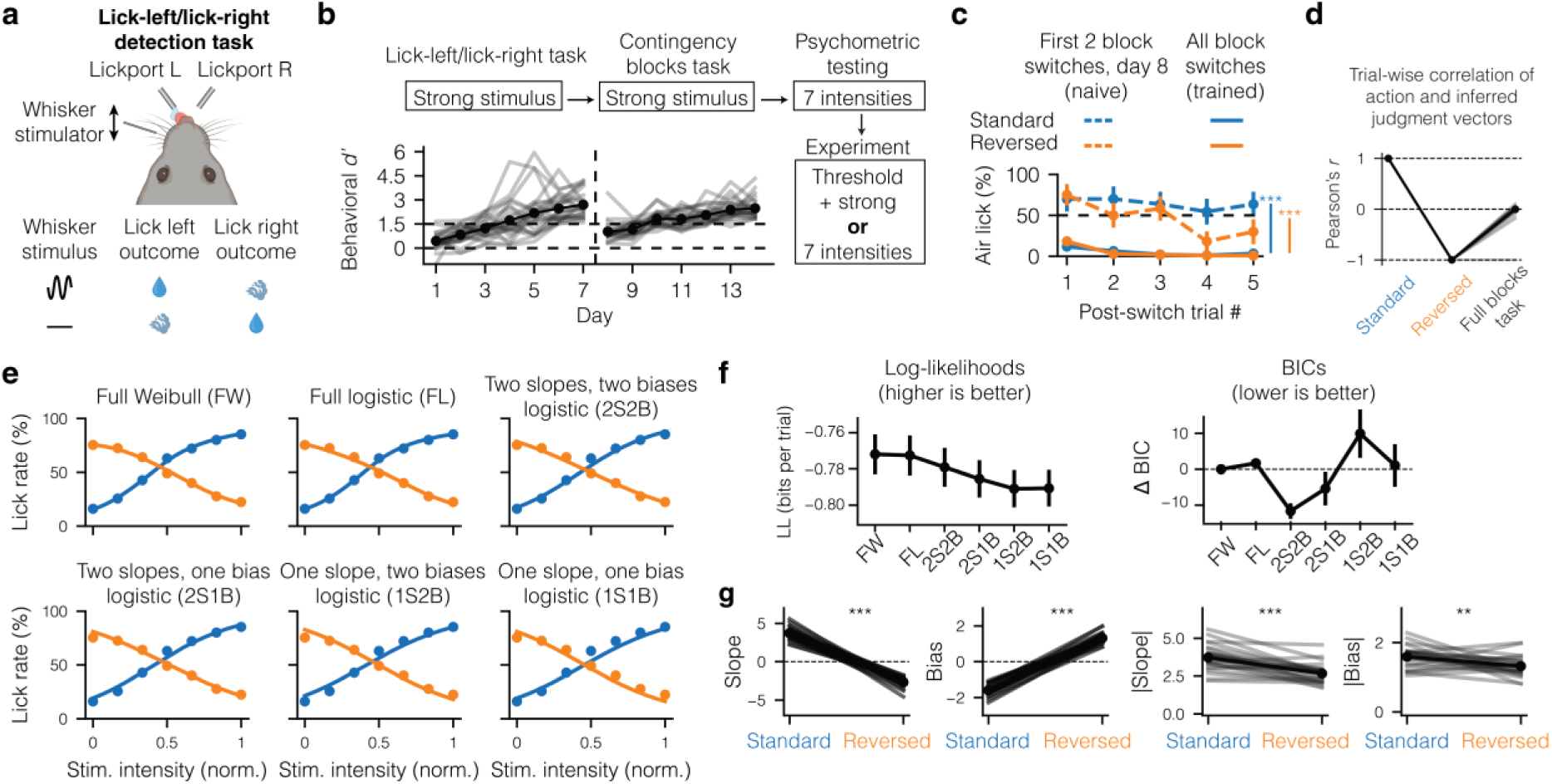
Additional task and learning data. (a) Schematic of the mouse performing the lick-left/lick-right detection task (top) and the contingency table of this task (bottom). (b) Learning curves across the lick-left/lick-right (pink) and blocks (blue) phases of task learning, as well as a schematic of all the experimental stages (top and right). (c) Air lick rate as a function of the number of post-switch trials, for either the first two block switches (day 8, naive, dotted lines) or all block switches in the experiment stage (trained, solid lines). Note the first post-switch trial was always a clear no-go trial, unlike the subsequent trials. We compared percent air licks and correct responses between the means of all post-switch trials across naive and trained sessions. (d) Pearson’s correlation between the action and inferred judgment vectors across only standard trials, only reversed trials, or both (*n* = 25 mice). Combining the two blocks breaks the otherwise definitional collinearity between the two. (e) Comparisons of six different models for fitting the psychometric data. For simplicity, only the overall fit on mouse-averaged data is shown. (f) Results of across-mouse log-likelihood (left) and Bayesian information criterion (BIC; right) calculations for each model. The two slopes, two biases (2S2B) model was most parsimonious based on yielding the lowest BIC. (g) Comparison of 2S2B parameters, with the first two panels from the left depicting the signed slopes and biases and the second two panels depicting their respective magnitudes. Significances in (c) and (g) were calculated using the two-sided Wilcoxon signed-rank test. * *P <* 0.05, ** *P <* 0.01, *** *P <* 0.001. Mouse illustration in (a) was partially created with BioRender.com.

**Extended Data Figure 2:**
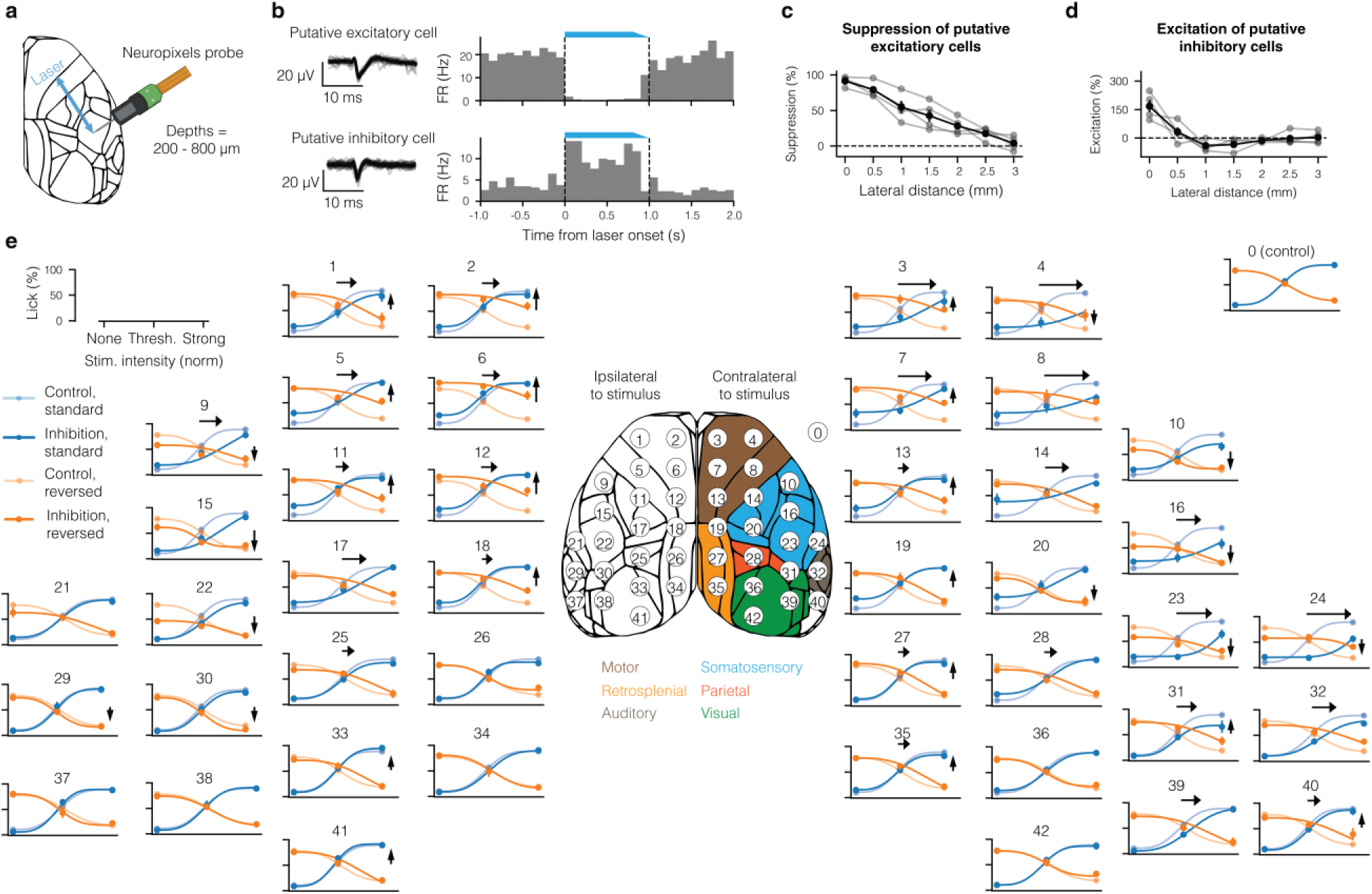
Detailed physiological and behavioral effects of inhibition. (a) A Neuropixels probe was inserted into S1-bc while optogenetic stimuli were given at distances between 0 and 3 mm in 0.5 mm steps. (b) Examples of a putative excitatory unit (top) and a putative inhibitory unit (bottom), each with its peri-stimulus time histogram during inhibition. (c) Suppression of putative excitatory units as a function of lateral distance between the laser and the insertion site for *n* = 4 mice. (d) As in (c) but for the excitation of putative inhibitory units. (e) Curves from all dorsal cortical regions across the standard and reversed blocks (see the legend at the top left), comparing the effect of inhibition targeted at each numbered region to the control curves (top right). Arrows are displayed only for effects greater than 5%. Error bars in (c), (d), and (e) show mean ± s.e.m. across *n* = 4 mice in (c) and (d) and *n* = 5 mice in (e). Mouse illustration in (a) was partially created with BioRender.com.

**Extended Data Figure 3:**
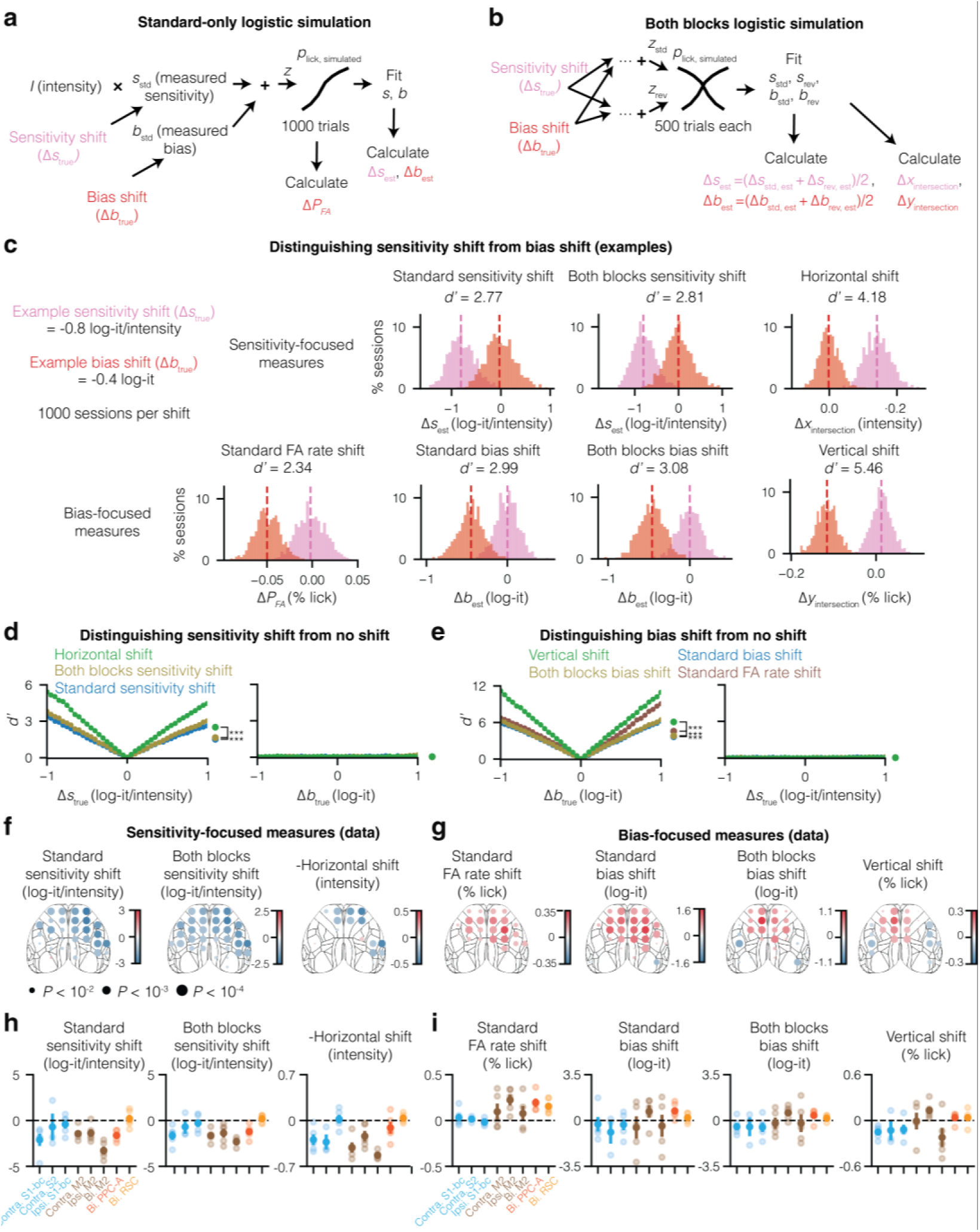
Intersection method best dissociates causal effects on sensitivity vs. bias. (a) Schematic showing how data is generated from a standard-only logistic simulation, followed by parameter estimation. FA: false alarm. (b) Same as (a) but both blocks simulation. (c) Comparison of shift estimation measures. The top row contains sensitivity-focused measures while the bottom contains bias-focused measures. In each case, the measure is compared across slope shift (pink) and bias shift (red) sessions. (d) A *d’* comparison of the sensitivity-focused approaches across various shift magnitudes. (e) Same as (d) but for bias-focused approaches. (f) Effects of inhibition in each screened region on the sensitivity-focused measures. Horizontal shifts are negated for comparison. (g) Same as (f) but for bias-focused approaches. (h) Effect of inhibition in each targeted region on the sensitivity-focused measures. Horizontal shifts are negated for comparison. (i) Same as (h) but for bias-focused approaches. Filled circles and error bars in (h) and (i) show mean ± s.e.m. across *n* = 5 mice, while unfilled circles show individual mice. *** *P <* 0.001

**Extended Data Figure 4:**
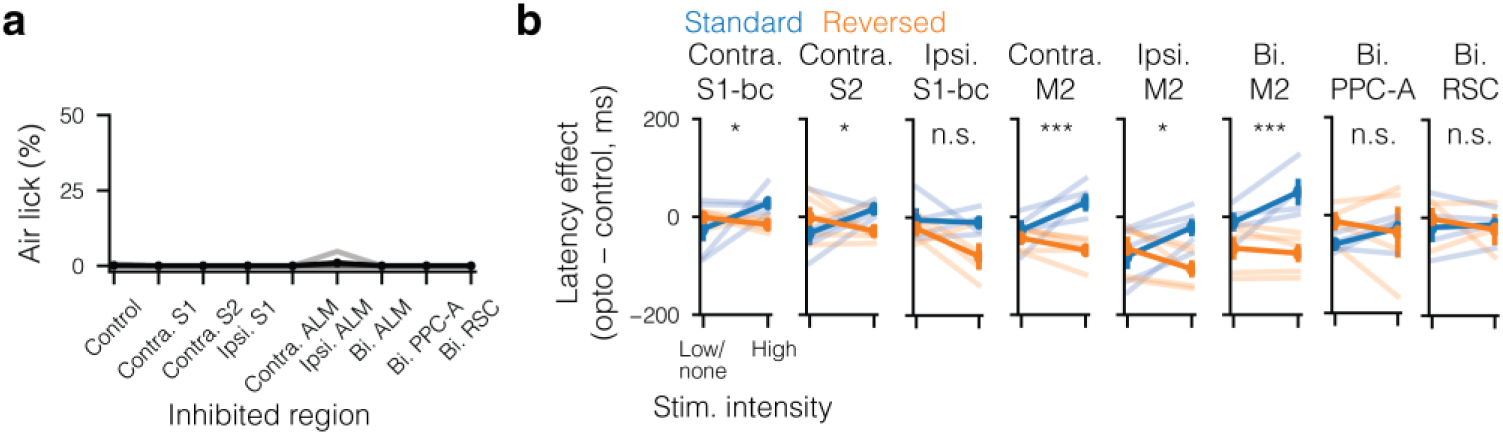
Effect of optoinhibition on air licks and lick latencies. (a) Percent air licks for each inhibition condition in the targeted inhibition experiment. No significant effects were found. (b) Effects of inhibition on lick latency across low (below threshold) and high (above threshold) stimulus intensities. Significance of each intensity × block interaction term was calculated using the Scheirer–Ray–Hare test, corrected for multiple comparisons using the Dunn–Šidák correction. Error bars show mean ± s.e.m. across *n* = 5 mice. * *P <* 0.05, ** *P <* 0.01, *** *P <* 0.001

**Extended Data Figure 5:**
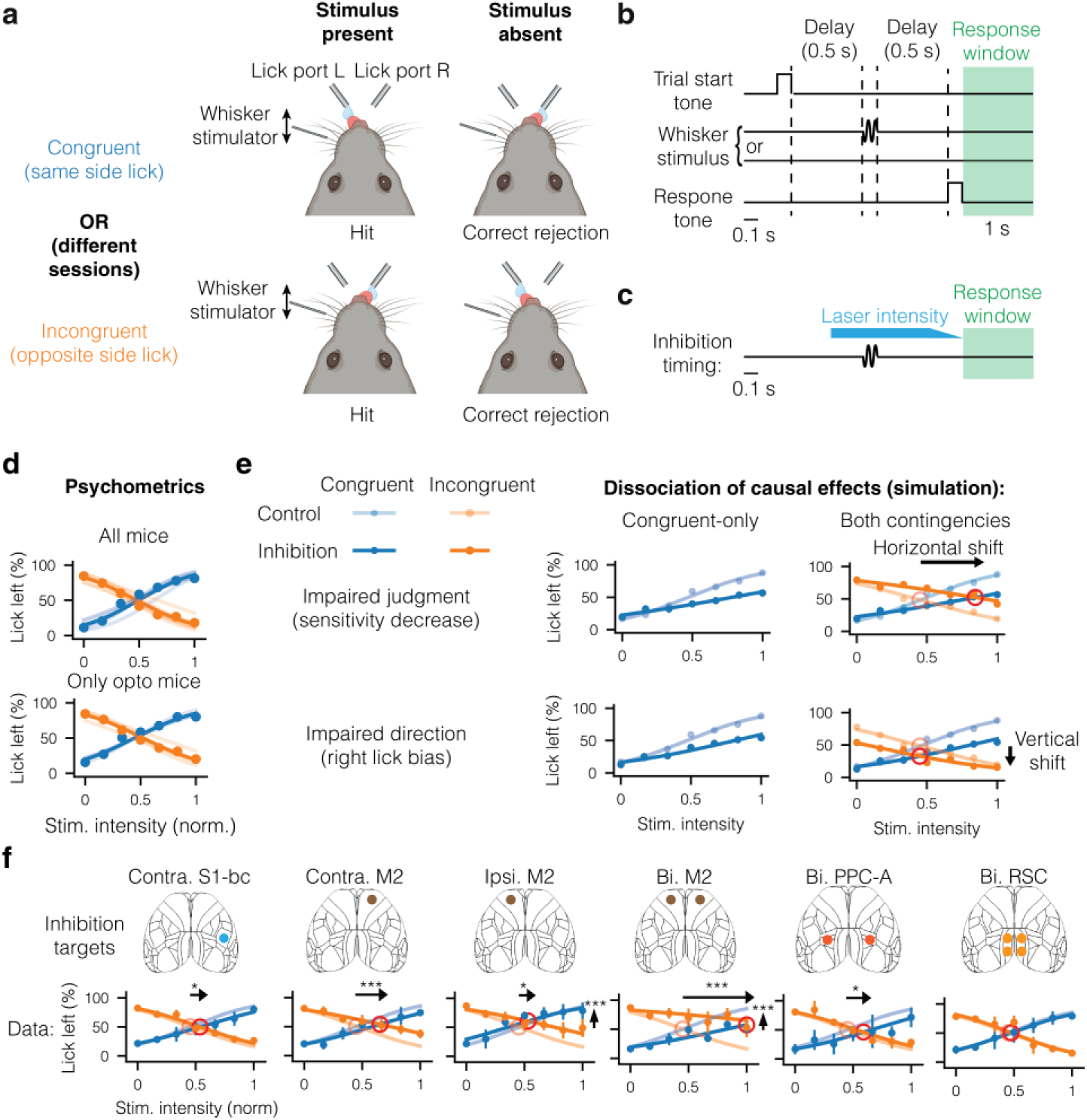
Optoinhibition experiment in the lick-left/lick-right task with delay. (a) Schematic of a mouse performing correct trials across stimulus present (left) and stimulus absent (right) trials on either congruent contingency (top) or reversed contingency (bottom). In the two mice that were trained on both contingencies, they were switched once for each mouse and required re-training. (b) Schematic of the trial structure in both contingencies. (c) Schematic of inhibition timing during both contingencies of the task. (d) Psychometric curves of left licks across contingencies for all mice trained on the lick-left/lick-right task (top) and only mice used for the optogenetics experiment (bottom). (e) Simulations of the effect of a decrease in sensitivity (top row) or an increase in right lick bias (bottom row) on left lick rate in the congruent contingency (left-hand column) or both contingencies (right-hand column). (f) Measured effect of inhibition in both contingencies on left lick rate. Error bars show mean ± s.e.m. across *n* = 3 mice in each contingency. * *P <* 0.05, ** *P <* 0.01, *** *P <* 0.001.

**Extended Data Figure 6:**
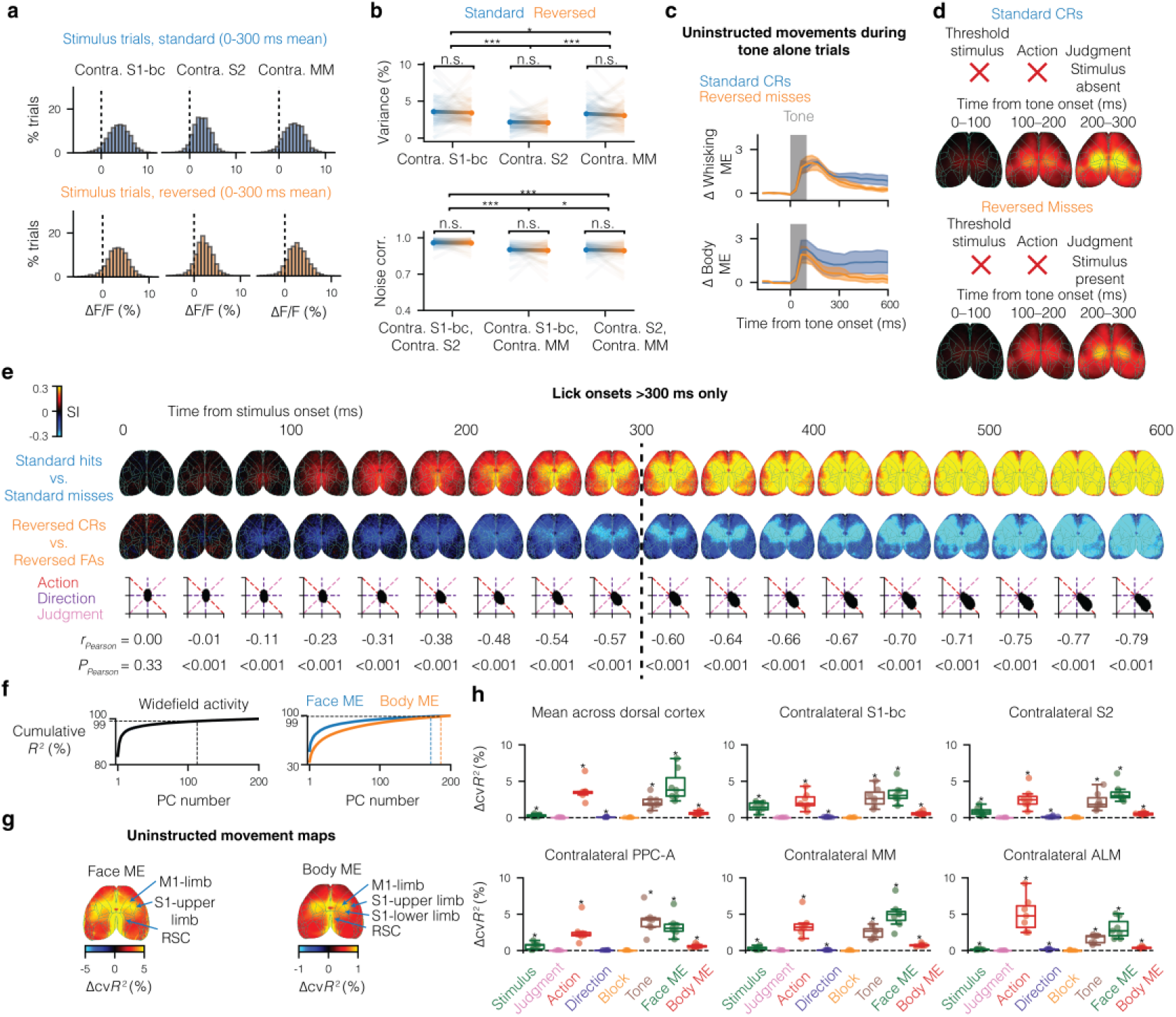
Additional analysis of widefield data. (a) Plots of single-trial mean responses between 0 and 300 ms after the stimulus across different regions (columns), namely contra. S1-bc (contralateral barrel cortex), contra. S2 (contralateral secondary somatosensory cortex), and contra. MM (contralateral medial motor cortex), and the two contingency blocks (rows). (b) Comparison of variances across regions (top) and comparison of noise correlations across region pairs (bottom). Transparent lines in the background are from the *n* = 72 sessions included, while the thick lines are their means with error bars indicating the s.e.m. across sessions. ANOVA with Šídák post-hoc tests across regions and contingency blocks. (c) In response to the tone, we observed an increase in whisking (left) and body movements (right) in both standard CRs and reversed misses. Error bands show mean ± s.e.m. across *n* = 7 mice. (d) Activity in both standard CRs and reversed misses displayed a global, bilateral pattern, which was concomitant with or closely followed the uninstructed movements in (c). (e) Time-lapse of comparisons between standard hits vs. misses and reversed CRs vs. FAs, as in Fig. 3f–h but restricted to trials in which mice licked after 300 ms and on a frame-by-frame basis. (f) Cumulative explained variance of widefield video data (left) and face and body motion energy video data (right) as a function of the number of PCs included for each. (g) Unique explained variances corresponding to uninstructed movements of the face (left) and body (right), with labels indicating peak regions. Note the difference in scales. (h) Region-averaged explained variances across mice. The boxes extend from the first quartile (Q1) to the third quartile (Q3) of the data, with a line at the median. The whiskers extend from the box to the farthest data point lying within 1.5× the interquartile range from the box. Significances were assessed using the one-sided Wilcoxon signed-rank test, corrected for multiple comparisons using the Dunn–Šidák correction. * *P <* 0.05, ** *P <* 0.01, *** *P <* 0.001.

**Extended Data Figure 7:**
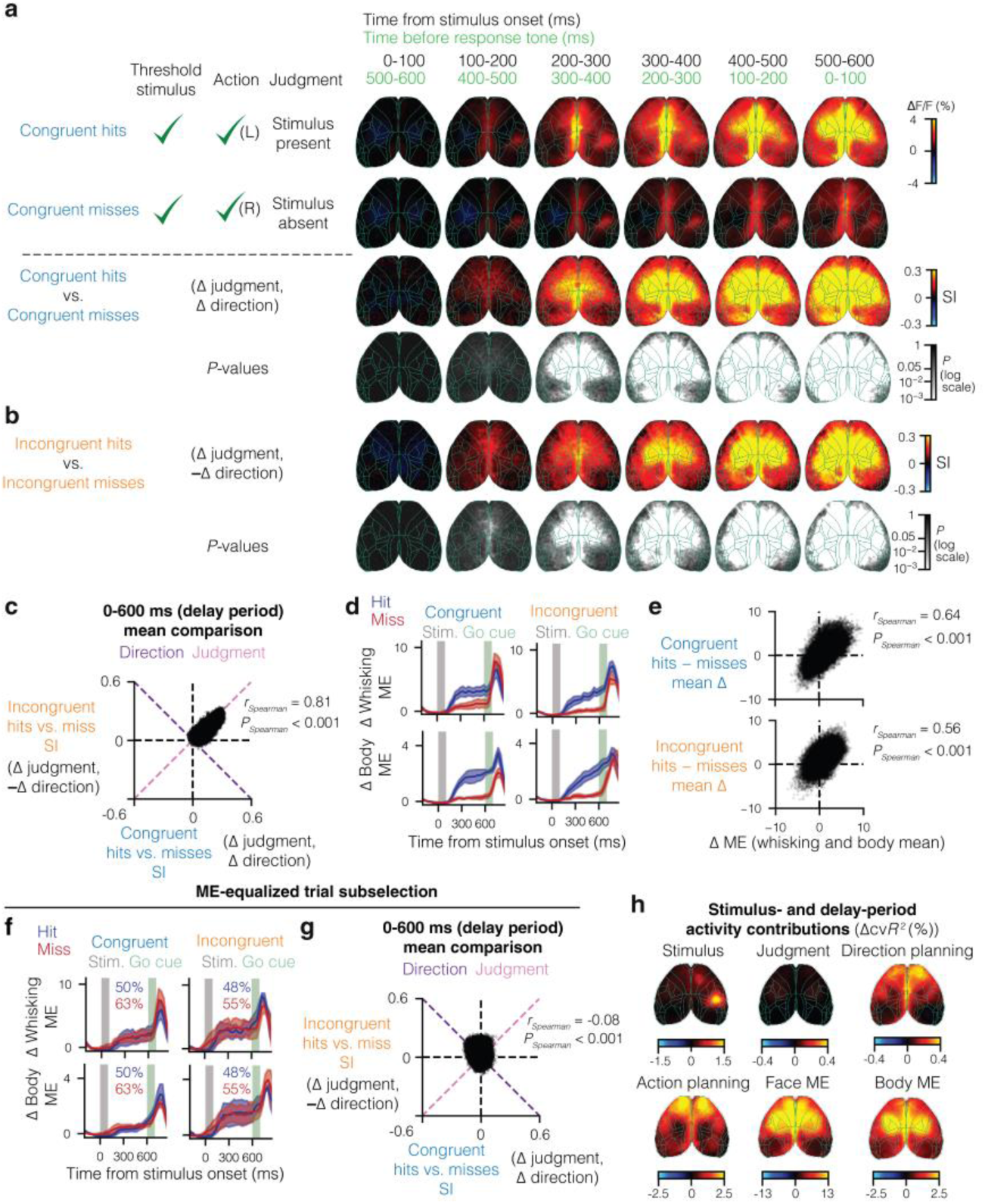
Widefield imaging experiments in the lick-left/lick-right task with delay. (a) Comparison of stimulus present vs. absent judgment trials in the congruent contingency of the lick-left/lick-right task. Top two rows: grand-averaged activity in hit (correct left lick) trials and in miss (incorrect right lick) trials. Bottom two rows: the results of the SI calculation between the two trial types and the Benjamini–Hochberg-corrected *P* values of the SIs obtained by comparing them to shuffled results. The columns on the left lay out the three factors defining each trial type (stimulus, action, and judgment), while the columns on the right represent the time bins from stimulus onset to the response tone. (b) Same as the bottom two rows of (a) but for the incongruent contingency. (c) Scatter plot of all widefield pixels across the comparisons in (a) and (b) revealed a positive relationship on a pixel-wise basis. Labels at the top are colored like the axes and diagonals along which lick direction-coupled and action-independent perceptual judgment correlates are expected to reside. (d) In contrast to the blocks task (Fig. 1f), uninstructed whisking (top) and body (bottom) movements were higher on stimulus present judgment trials (hits) compared to stimulus absent judgment trials (misses) in both contingencies. (e) Scatter plot of all pairwise differences in mean cortex-wide activity compared to pairwise differences in mean of uninstructed movements (whisking and body) across the delay period. The two were positively correlated for both congruent (top) and incongruent (bottom) contingencies. (f) Same as (d) but for a subselection of hits and misses designed to equalize the mean uninstructed motion energies (MEs) across trials. Percents inside each subpanel indicate the percent of hit (blue) and miss trials (red) that were included. (g) Same as (c) but among subselected trials only. (h) Unique explained variances for delay-period predictors from a ridge regression applied to the full dataset. Error bands in (d) and (f) show mean ± s.e.m. across *n* = 4 mice in each contingency.

**Extended Data Figure 8:**
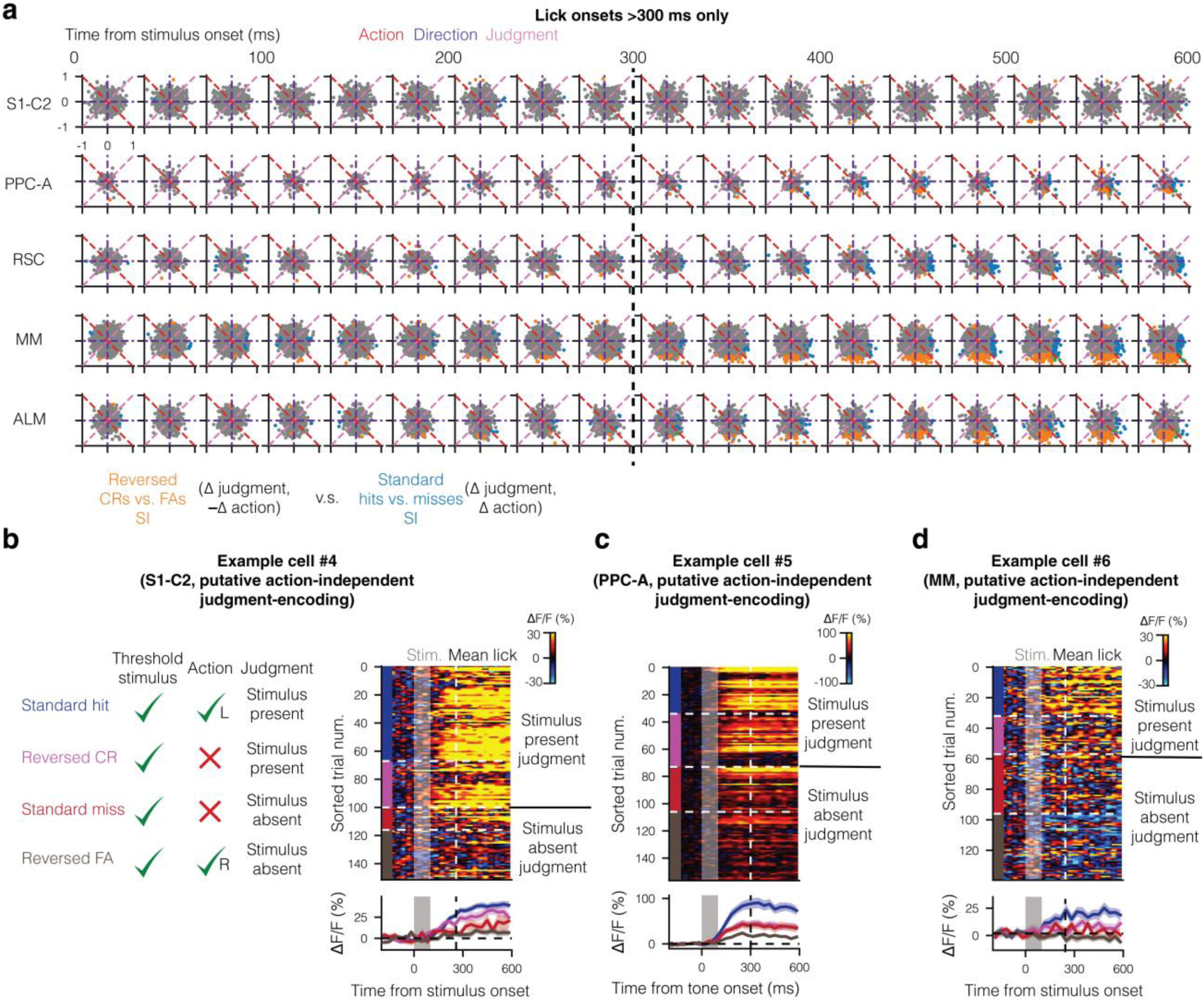
Deeper search for single-cell perceptual judgment correlates. (a) Time-lapse of reversed CRs vs. FAs SI (abscissa) plotted against standard hits vs. misses SI (ordinate), as in Fig. 4h but only for trials in which mice licked past 300 ms and on a frame-by-frame basis. (b) An example of a putative S1-C2 action-independent judgment-encoding cell, with trial labels on the left, raster plot at the top-right, and peri-stimulus time histogram (PSTH) at the bottom-right. (c,d) As in (b), but for a putative PPC-A action-independent judgment-encoding cell and an MM action-independent judgment cell, respectively. Error bands in PSTHs in (b-d) show mean ± s.e.m. across fields of view.

**Extended Data Figure 9:**
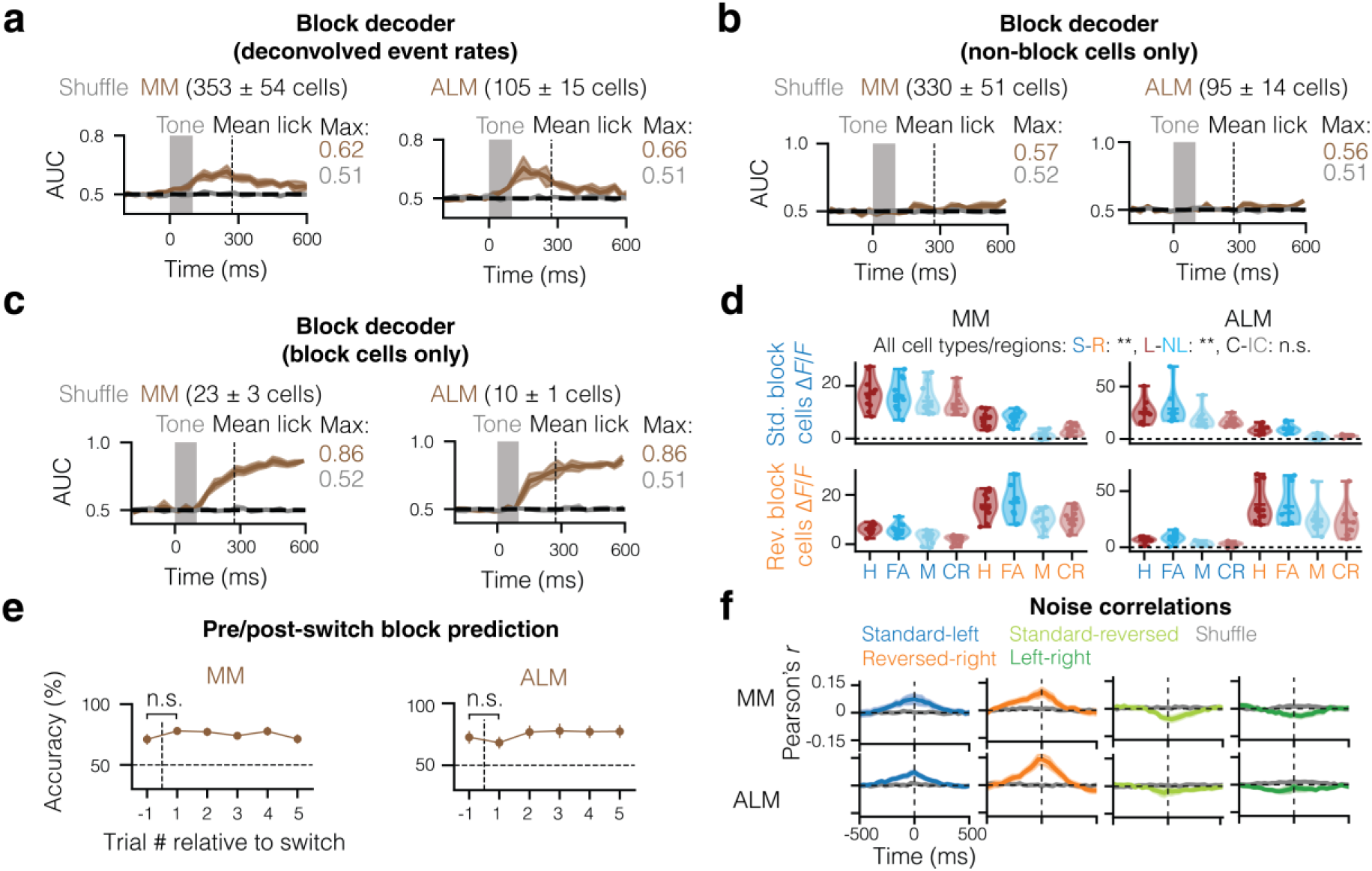
Additional exploration of block coding. (a) Result of block decoding, as in Fig. 5c but using deconvolved event rates rather than Δ*F/F*. (b) Result of block decoding, as in Fig. 5c but after removing cells that individually significantly encoded the block. Number of cells is mean ± s.e.m. across mice. (c) Same as (b) but after removing cells that did not individually significantly encode the block, i.e., classification based on block cells alone. Number of cells is mean ± s.e.m. across mice. (d) Comparison of standard and reversed block cell mean Δ*F/F* (0-600 ms post-stimulus) across different regions and trial types. H: hits, FA: false alarms, M: misses, CR: correct rejection. S-R is the comparison across standard and reversed blocks, L-NL is the comparison across lick and no lick trials, and C-IC is the comparison across correct and incorrect trials. All comparisons were performed separately for each block cell population and region (corrected for multiple comparisons with the Dunn–Šidák correction), but the statistical results were the same. (e) Accuracy of block prediction using the population activity (Δ*F/F*) across different trials relative to each switch, for both MM (left) and ALM (right). (f) Noise correlations in MM (upper row) and ALM (lower row) between, from left to right, standard block and lick left cells, reversed block and lick right cells, standard and reversed block cells, and lick left and lick right cells. Error bands show mean ± s.e.m. across FOVs from each region. * *P <* 0.05, ** *P <* 0.01, *** *P <* 0.001. n.s. (not significant) indicates *P* > 0.05.

**Extended Data Figure 10:**
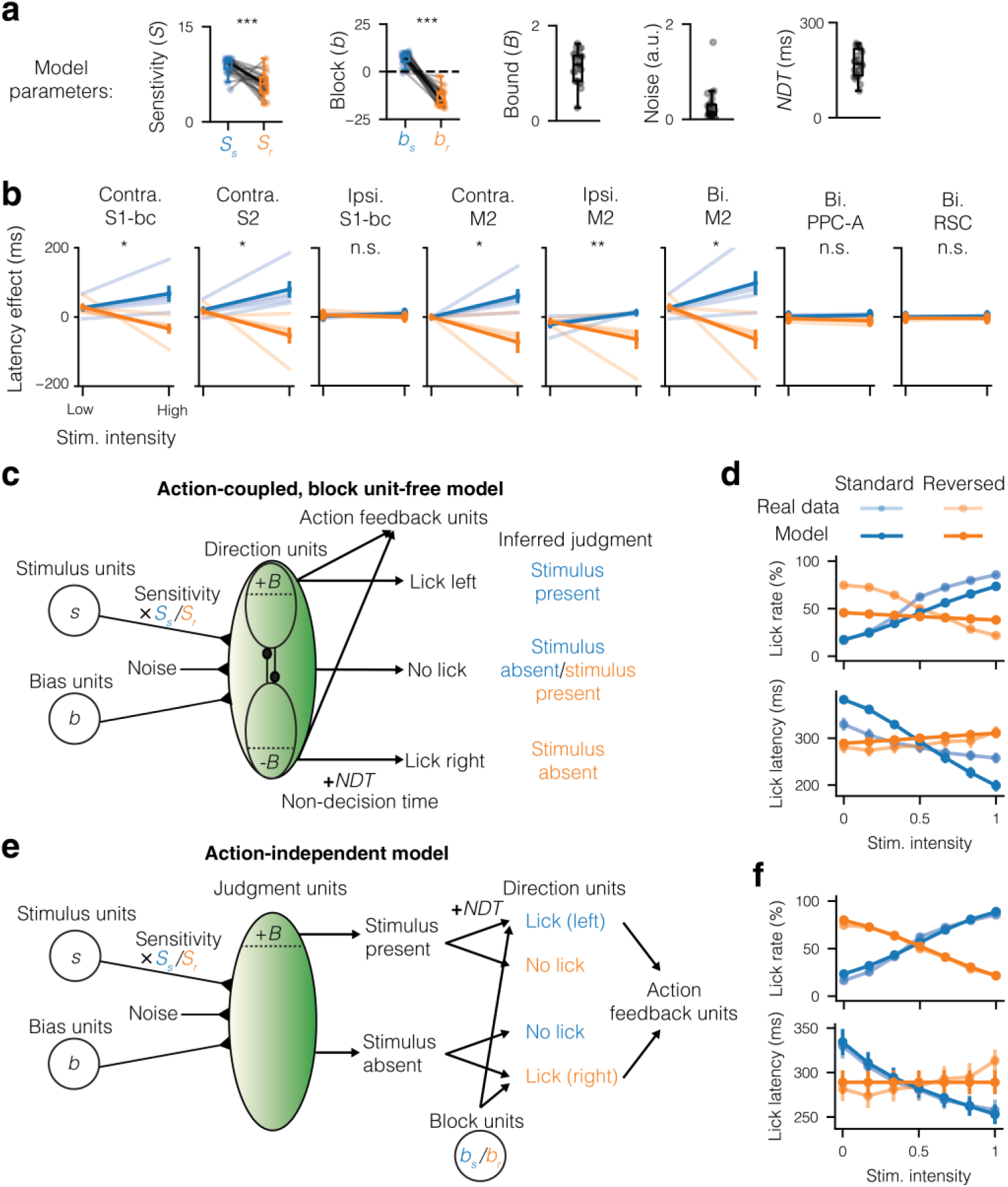
Additional model results and alternative models. (a) Final parameters of the fitted action-coupled model in Fig. 6, including a cross-block comparison of the sensitivity parameter and block (bias) input. Significances were assessed using the two-sided Wilcoxon signed-rank test. (b) Lick latency effect of shifting model parameters to fit the optogenetic inhibition. Significance of each intensity × block interaction term was calculated using the Scheirer–Ray–Hare test, corrected for multiple comparisons using the Dunn–Šidák correction. Compare to Extended Data Fig. 4b. (c) Action-coupled model but with the block units replaced by nonspecific bias units. (d) Results of fitting the block unit-free model to the data with the same approach as in Fig. 6c. (e) An action-independent judgment model. In contrast to the other models, judgment is represented explicitly in this model. (f) Results of fitting the action-independent judgment model to the data with the same approach as in Fig. 6c. Error bars show mean ± s.e.m. across fits to individual mice. * *P <* 0.05, ** *P <* 0.01, *** *P <* 0.001.

