## Supplementary Figures 1-4 and Supplementary Note 1 for "Inherent coupling of perceptual judgments to actions in the mouse cortex"

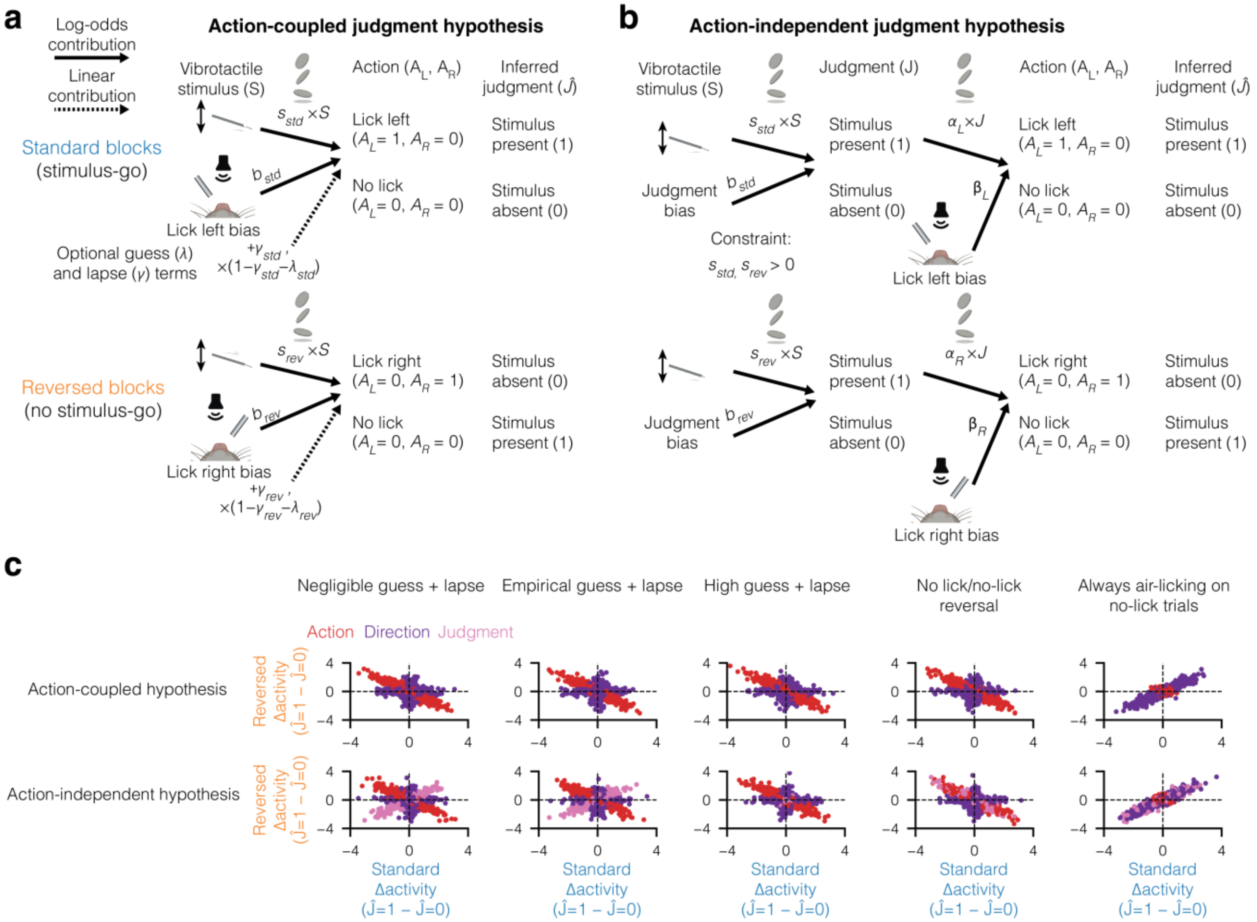


**Supplementary Figure 1: Neural separation of judgment and action in the blocks task. (a)** Schema of the contribution of stimulus, bias, guess, and lapse parameters to the probability of licking left vs. not licking in standard blocks (top) and licking right vs. not licking in the reversed blocks (bottom). This schema assumes the action-coupled judgment hypothesis, which posits there is no intermediate stage in which a judgment is computed prior to the action. **(b)** Same as (a) but for the action-independent judgment hypothesis, which posits there is an intermediate stage in which judgment is computed prior to action. **(c)** Simulated neural activity differences across within-block judgment contrasts for the action-coupled judgment hypothesis (top row) and action-independent judgment hypothesis (bottom row). Each column represents different model parameter settings, except the last column which fully replaces no-lick trials with air licks; see Supplementary Note 1 for details. Dot colors represent the type of neural variable, with non-specific action and uninstructed movement variables in red, left and right direction variables in purple, and judgment variables in pink.

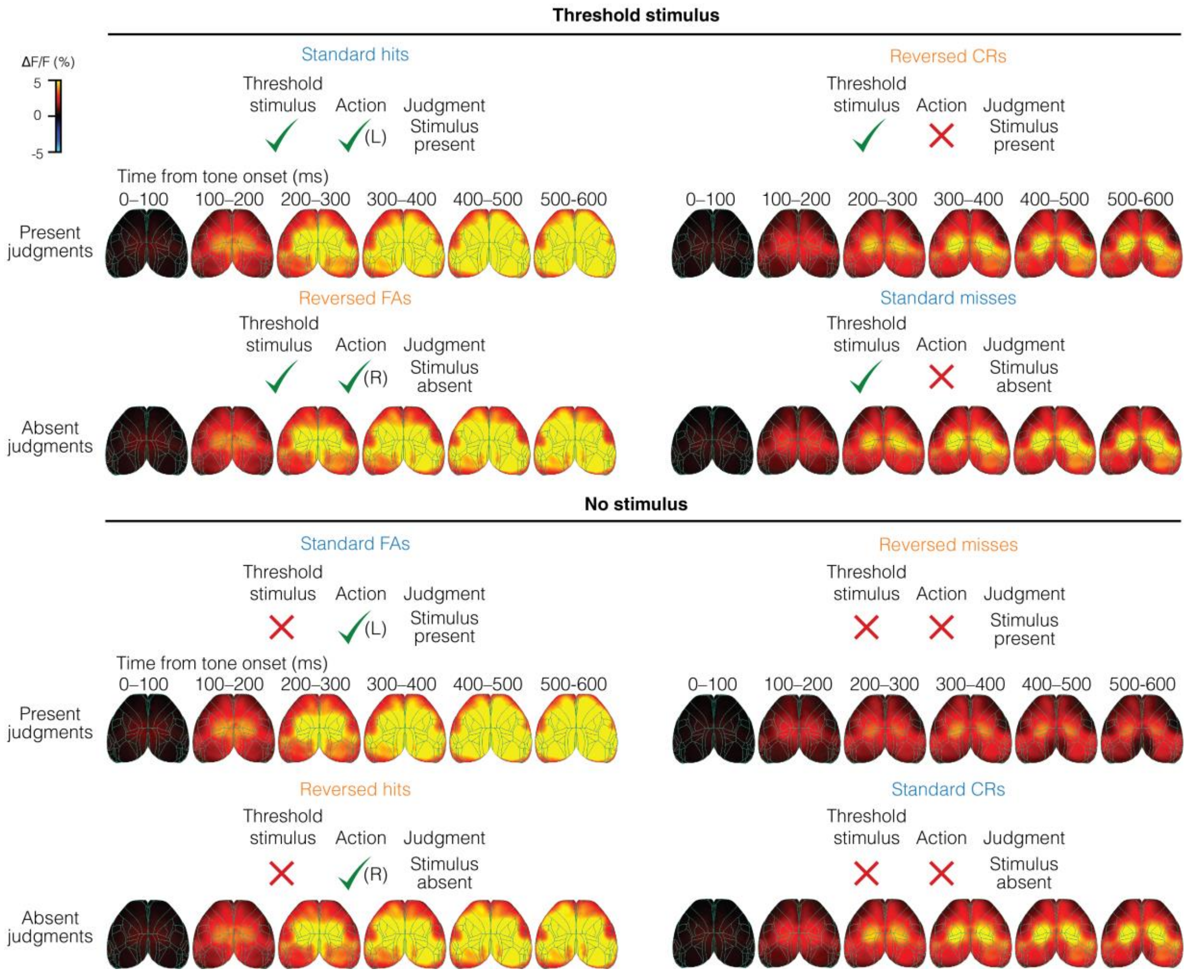


**Supplementary Figure 2: Stimulus- and action-matched widefield comparisons of all trial types across blocks.** Matrix of widefield responses across trial types, arranged by threshold stimulus vs. no stimulus (top vs. bottom half), lick vs. no lick (left vs. right half), and present vs. absent judgment (upper vs. lower rows in each stimulus condition).

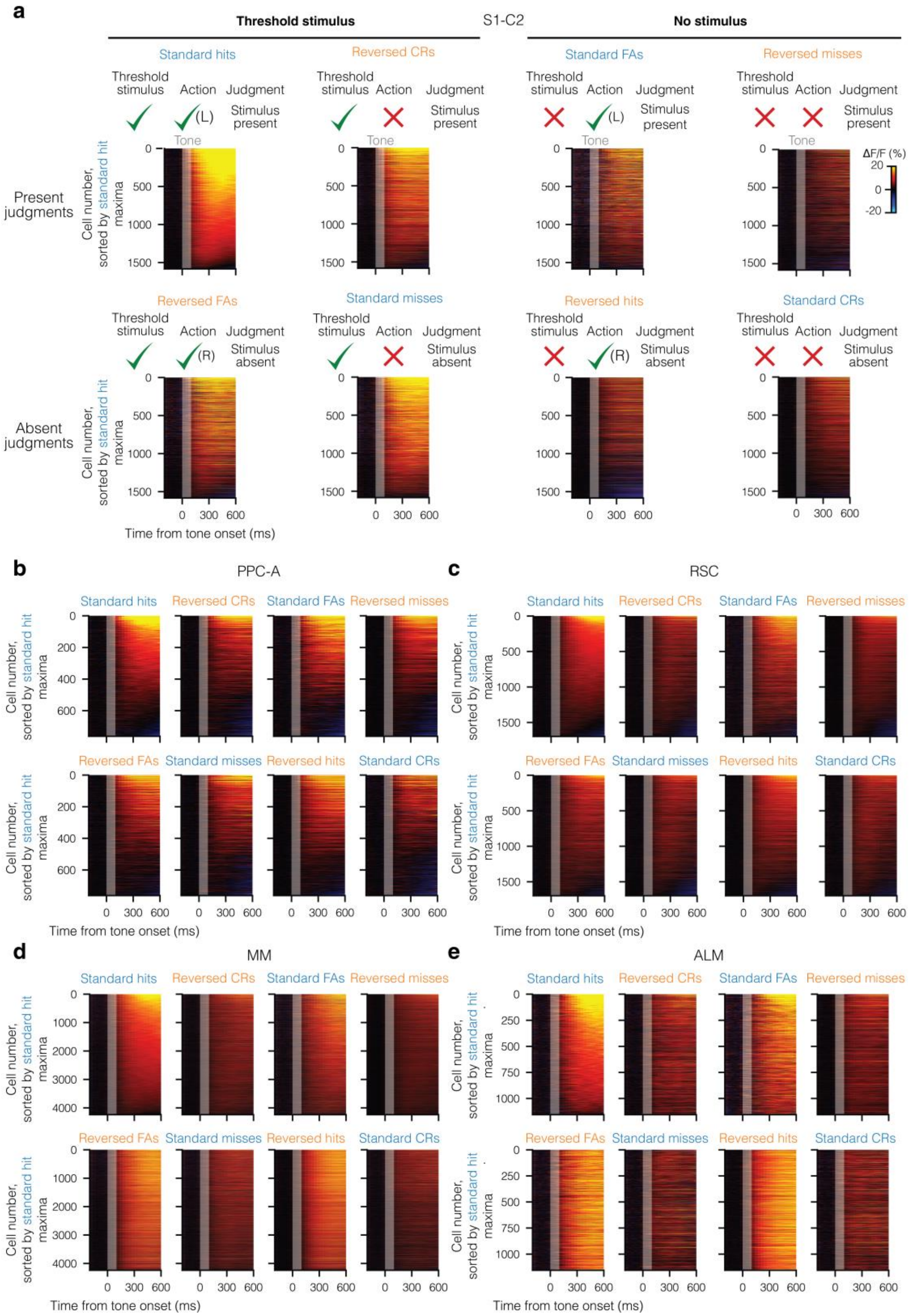


**Supplementary Figure 3: Stimulus- and action-matched two-photon comparisons of all trial types across blocks. (a)** All recorded S1-C2 cells, sorted according to mean activity on standard hit trials (top-left corner). Trial types arranged as in Supplementary Fig. 2. **(b-e)** Same as (a), but with all recorded PPC-A, RSC, MM, and ALM cells, respectively.

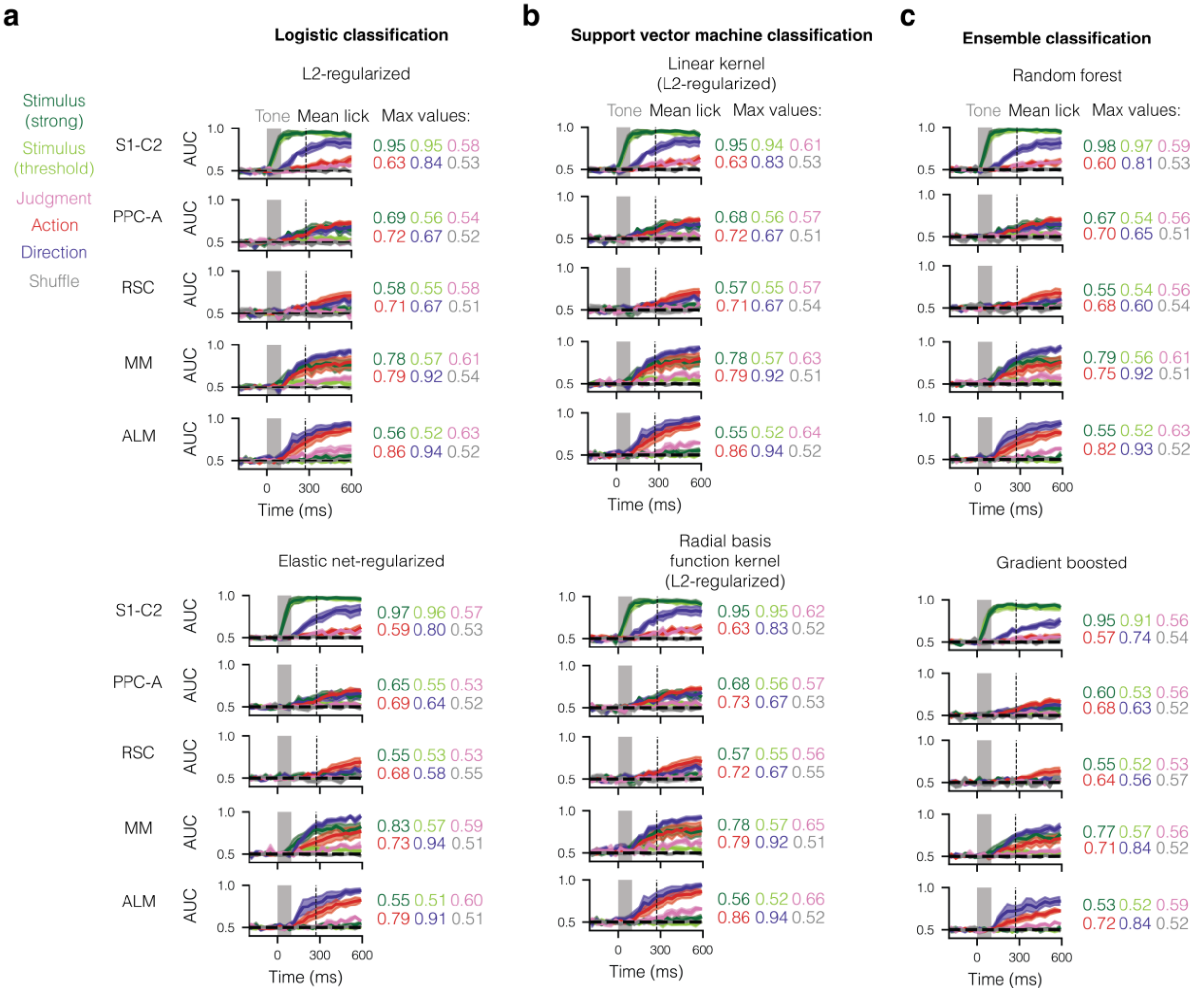


**Supplementary Figure 4: Different classifiers reveal similar factor magnitudes and time courses. (a)** Same as in Fig. 4j but with L2-regularized (top) or elastic net-regularized (bottom) logistic classifiers. **(b)** Same but with linear kernel (top) or radial basis function kernel (bottom) support vector machine classifiers. **(c)** Same but with nonlinear random forest (top) or gradient-boosted (bottom) decision tree classifiers. Error bands show mean ± s.e.m. across fields of view.

### Supplementary Note 1: Judgment-Action Dissociation Via Within-Block Comparisons

In this note, we consider two hypotheses about how perceptual judgments are made: action-independent and action-coupled. We show that under common assumptions, the two are not distinguishable in terms of behavior alone but can be distinguished neurally. In particular, we derive the assumptions underlying the central claim that it is possible to neurally decouple perceptual judgments from actions in the blocks task (Fig. 1) if the action-independent judgment hypothesis is correct.

Let us express the evidence to lick left ($z_L$) or right ($z_R$) under the **action-coupled judgment hypothesis** (Supplementary Fig. 1a), in which each probability is directly influenced by the whisker stimulus input ($S$) and lick direction biases ($b_L$ and $b_R$). In addition, to account for behavior change upon block switches, we add a term $A(L)/A(R)$ signifying the absence of a lick port on the same side (or, equivalently, the presence of a lick port on the other side):

$$z_L = s_L S - a_L A(L) + b_L \quad (1)$$
$$z_R = s_R S - a_R A(R) + b_R \quad (2)$$

where $s_L$ and $s_R$ are the whisker stimulus weights and $a_L$ and $a_R$ are the lick port absence weights. As air licks are very rare (Extended Data Fig. 1c) and are excluded from analysis, this causes the $a$ parameters to diverge. Hence, we can only model $z_L$ on standard blocks (where $A(L) = 0$) and $z_R$ on reversed blocks (where $A(R) = 0$):

$$z_{L|std} = s_{std} S + b_{std} \quad (3)$$
$$z_{R|rev} = s_{rev} S + b_{rev} \quad (4)$$

We incorporate this evidence to the lick probability on each block using a logistic function, alongside two possible evidence-independent terms, namely a guess rate (γ; lower asymptote) and a lapse rate (λ; 1-higher asymptote), which may be different across blocks:

$$P(L|std) = \gamma_{std} + (1 - \gamma_{std} - \lambda_{std}) \times \sigma(z_{L|std}) \quad (5)$$
$$P(R|rev) = \gamma_{rev} + (1 - \gamma_{rev} - \lambda_{rev}) \times \sigma(z_{R|rev}) \quad (6)$$

where $\sigma$ is the sigmoid function ($\sigma(x) = 1/(1 + e^{-x})$).

Under the **action-independent judgment hypothesis** (Supplementary Fig. 1b), the evidence is expressed in the same manner as in equations (3) and (4), but rather than directly contributing to the probability of licking left or right on each block, it contributes to an intermediate probability of judging the stimulus as present ($P(J)$):

$$P(J|std) = \sigma(z_{std}) \quad (7)$$
$$P(J|rev) = \sigma(z_{rev}) \quad (8)$$

An important constraint of this hypothesis but not the action-coupled hypothesis is that both $s_{std}$ and $s_{rev}$ are > 0, which ensures that the resulting judgment probability goes up monotonically with stimulus strength. The lick probability is then linked to this judgment via another set of equations:

$$P(L|std) = \sigma(\alpha_L J + \beta_L) \quad (9)$$
$$P(R|rev) = \sigma(\alpha_R J + \beta_R) \quad (10)$$

where $\alpha_L$ and $\alpha_R$ are the weights of stimulus presence judgments towards licking left and right, and $\beta_L$ and $\beta_R$ are the direction biases. While we could additionally add guess and lapse terms to either set of equations, we will argue that doing so is superfluous because the $\alpha$ and $\beta$ parameters themselves account for guesses and lapses in the classical sense. Specifically, guesses are classically defined as the probability of a specific response when the evidence *towards* the response is absent. In the standard block this is simply

$$\gamma_{std} = P(L|std, \sigma(z_{std}) = 0) = \sigma(\beta_L) \tag{11}$$

However, in the reversed block the evidence $z_{R|rev}$, which increases monotonically with P(J), is *against* the lick right response. Thus guesses occur when its complement is zero:

$$\gamma_{rev} = P(R|rev, 1 - \sigma(z_{rev}) = 0) = \sigma(\alpha_R + \beta_R) \tag{12}$$

On the other hand, lapses are classically defined as the probability of failing to make a specific response when the evidence towards the response is maximal, so by the same logic:

$$\lambda_{std} = 1 - P(L|std, \sigma(z_{std}) = 1) = 1 - \sigma(\alpha_L + \beta_L) \tag{13}$$
$$\lambda_{rev} = 1 - P(R|rev, 1 - \sigma(z_{rev}) = 1) = 1 - \sigma(\beta_R) \tag{14}$$

To achieve the particular case of $\gamma_{std} = 0$ and $\lambda_{std} = 0$, $\beta_L$ would thus need to be $\ll 0$ while $\alpha_L$ would need to be both $\gg 0$ and $\gg \beta_L$ in magnitude. Meanwhile, to achieve $\gamma_{rev} = 0$ and $\lambda_{rev} = 0$, $\beta_R$ would need to be $\gg 0$ while $\alpha_R$ would need to be both $\ll 0$ and $\gg \beta_R$ in magnitude. The result of this parameter choice is a perfect, step-function second stage that always produces the correct action in each contingency given the judgment.

Do the two hypotheses make different behavioral predictions? While the double-sigmoid form of the action-independent equations cannot be reduced to a linear operation on a single sigmoid as in the action-coupled hypothesis, in the above limit of very low guess and lapse rates we have:

$$P(L|std) \approx P(J|std) = \sigma(z_{std}) = \sigma(s_{std}S + b_{std}) \tag{15}$$
$$P(R|rev) \approx 1 - P(J|rev) = 1 - \sigma(z_{rev}) = 1 - \sigma(s_{rev}S + b_{rev}) = \sigma(-s_{rev}S - b_{rev}) \tag{16}$$

Therefore, in this limit the two hypotheses converge to make the same behavioral predictions. While in standard blocks this happens with identical sensitivity and bias parameters, in reversed blocks it happens with *opposite* sensitivity and bias parameters. The constraint of positive sensitivities in the action-independent hypothesis thus implies that the sensitivities need to have opposite signs across blocks in the action-coupled hypothesis, namely positive for standard and negative for reversed, as empirically observed (Extended Data Fig. 1g).

Next, we consider whether it is possible to distinguish between the two hypotheses based on neural activity. Let $X_i$ represent a neural variable (pixel activity, single neuron response, etc.), and let it be a linear combination of different task variables:

$$X_i = w_{T,i}T + w_{S,i}S + w_{Pr_L,i}Pr_L + w_{Pr_R,i}Pr_R + w_{J,i}J + w_{A_L,i}A_L + w_{A_R,i}A_R + w_{A,i} * \{A_L = 1|A_R = 1\} + w_{UI,i}\,UI \tag{17}$$

where $T$ represents the tone, $S$ represents the vibrotactile stimulus, $Pr_L$ and $Pr_R$ represent the left and right lick port presence (or absence of the other lick port), J represents the judgment, $A_L$ and $A_R$ represent left and right licks, and $UI$ represents the magnitude of uninstructed movements. The $w$s are the respective weights of each factor, which may be zero or non-zero. Specifically, in the case of the action-coupled hypothesis, $w_{J,i}$ is zero for every $X_i$ (as there is no intermediate computation of judgment) while in the case of the action-independent hypothesis, $w_{J,i}$ is non-zero for some $X_i$ (as there is an intermediate computation of judgment).

Now define the inferred judgment $\hat{J}$ as 1 on lick left trials in the standard block and no lick trials in the reversed blocks, and 0 otherwise, and the **within-block judgment contrast** in the standard and reversed blocks as:

$$\Delta_{std}X_i = E\left(X_i \middle| \hat{J} = 1, S = S_{thr}, std\right) - E\left(X_i \middle| \hat{J} = 0, S = S_{thr}, std\right) \tag{18}$$

$$\Delta_{rev}X_i = E\left(X_i \middle| \hat{J} = 1, S = S_{thr}, rev\right) - E\left(X_i \middle| \hat{J} = 0, S = S_{thr}, rev\right) \tag{19}$$

where $S_{thr}$ is the threshold stimulus intensity. The first contrast is between the expected value of activity in standard hits and misses, while the second is between the expected values of activity in reversed correct rejections and false alarms (as in Figs. 3h and 4h). Across the two contrasts, all factors which are identical (tone, stimulus, lick port presence) cancel out, leaving:

$$\Delta_{std}X_i = w_{J,i} * \Delta_{std}E(J) + w_{A_L,i} + w_{A,i} + w_{UI,i} * \Delta_{std}E[UI] \tag{20}$$

$$\Delta_{rev}X_i = w_{J,i} * \Delta_{rev}E(J) - w_{A_R,i} - w_{A,i} + w_{UI,i} * \Delta_{rev}E[UI] \tag{21}$$

Before we can simulate populations with different *w*s, we must first calculate the value of **(1)** $E(J)$ and **(2)** $E[UI]$ in each trial type. We will treat them in turn:

**(1)** In the action-coupled hypothesis, $w_{J,i} = 0$ for all units and thus the $E(J)$ terms can be ignored. In the action-independent hypothesis, we can solve for them by applying Bayes' theorem. For example for $\Delta_{std}E(J)$:

$$\Delta_{std}E(J) = P\left(J = 1 \middle| \hat{J} = 1, S = S_{thr}, std\right) - P\left(J = 1 \middle| \hat{J} = 0, S = S_{thr}, std\right)$$

$$= \frac{P\left(\hat{J} = 1, S = S_{thr}, std \middle| J = 1\right) * P(J = 1 | S = S_{thr}, std)}{P\left(\hat{J} = 1, S = S_{thr}, std \middle| J = 1\right) * P(J = 1 | S = S_{thr}, std) + P\left(\hat{J} = 1, S = S_{thr}, std \middle| J = 0\right) * P(J = 0 | S = S_{thr}, std)}$$

$$- \frac{P\left(\hat{J} = 0, S = S_{thr}, std \middle| J = 1\right) * P(J = 1 | S = S_{thr}, std)}{P\left(\hat{J} = 0, S = S_{thr}, std \middle| J = 0\right) * P(J = 1 | S = S_{thr}, std) + P\left(\hat{J} = 0, S = S_{thr}, std \middle| J = 0\right) * P(J = 0 | S = S_{thr}, std)} \tag{22}$$

Given that $S = S_{thr}$, we can assume both $P(J = 1 | S = S_{thr}, std)$ and $P(J = 0 | S = S_{thr}, std)$ are approximately equal to 0.5, while from Eqs. (11) and (13) we have $P\left(\hat{J} = 1, S = S_{thr}, std \middle| J = 1\right) = 1 - \lambda_{std}$ and $P\left(\hat{J} = 1, S = S_{thr}, std \middle| J = 0\right) = \gamma_{std}$. Thus

$$\Delta_{std}E(J) = \frac{1 - \lambda_{std}}{(1 - \lambda_{std}) + \gamma_{std}} - \frac{\lambda_{std}}{\lambda_{std} + (1 - \gamma_{std})} = \frac{1 - \gamma_{std} - \lambda_{std}}{1 - (\lambda_{std} - \gamma_{std})^2} \tag{23}$$

Similarly, it can be shown that

$$\Delta_{rev}E(J) = \frac{1 - \gamma_{rev} - \lambda_{rev}}{1 - (\lambda_{rev} - \gamma_{rev})^2} \tag{24}$$

If both $\gamma$s and $\lambda$s are very low, then, $\Delta_{std}E(J) \approx \Delta_{rev}E(J) \approx 1$.

**(2)** The values of $E[UI]$ on each trial type are an empirical question, and depend greatly on how uninstructed movements are defined. If we define uninstructed movements by the total change in motion energies of the body, whisker pad, or both, we know empirically that $\Delta_{std}E[UI] = E[UI|\text{std hits}] - E[UI|\text{std misses}] > 0$ while $\Delta_{rev}E[UI] = E[UI|\text{rev CRs}] - E[UI|\text{rev FAs}] < 0$, with $\Delta_{std}E[UI]$ and $\Delta_{rev}E[UI]$ having similar magnitudes (Fig. 1f). Uninstructed movements may also refer to significant principal components of movement in face and body videos following orthogonalization with respect to licks, which we also observed to be greater on standard hits and reversed FAs (data not shown). Thus, for simplicity, we assume $\Delta_{std}E[UI] = 1$ and $\Delta_{rev}E[UI] = -1$. As the resulting uninstructed movement contribution has the same sign and magnitude across the contrasts as the action contribution, we will group the two together as 'action-related'.

As a result, we obtain the following set of equations for the action-coupled hypothesis:

$$\Delta_{std} X_i = w_{A_L,i} + w_{A,i} + w_{UI,i} \tag{25}$$
$$\Delta_{rev} X_i = -w_{A_R,i} - w_{A,i} - w_{UI,i} \tag{26}$$

And for the action-independent hypothesis:

$$\Delta_{std} X_i = w_{J,i} + w_{A_L,i} + w_{A,i} + w_{UI,i} \tag{27}$$
$$\Delta_{rev} X_i = w_{J,i} - w_{A_R,i} - w_{A,i} - w_{UI,i} \tag{28}$$

We can group the different parts of each contrast by their relationship across the contrasts. Judgment (pink) activity is expected to be positively correlated across the contrasts, direction-related (purple) activity is expected to be uncorrelated across the contrasts, and action-related (red) activity is expected to be negatively correlated across the contrasts. To make this more concrete, we simulated a variety of $X_i$ with pure variable coding randomly chosen from each of the categories (Gaussian weights, σ = 1), added Gaussian noise (σ = 0.2), and computed the within-block contrasts. In the left-most column of Supplementary Fig. 1c (negligible guess + lapse), we plot the results of this simulation. Here, it is possible to pick out positively correlated neural variables in the action-independent judgment model, which correspond to judgment coding. Thus, under the specific set of assumptions we used above, it should be possible to neurally separate significant judgment coding from both action-related and direction-related coding, and to prove or falsify the action-independent coding hypothesis in specific recorded regions and coding schemes.

Next, we would also like to know what specific assumptions used above contribute to the observed separability of judgment coding. We therefore consider different violations of these assumptions:

**(1)** Empirical guess + lapse. Here we apply the guess and lapse rate values estimated from the data (full logistic model, Extended Data Fig. 1e), which were $\gamma_{std}$ = 0.073, $\lambda_{std}$ = 0.040, $\gamma_{rev}$ = 0.030, $\lambda_{rev}$ = 0.107. Plugging these values into the $\Delta_{std} E(J)$ and $\Delta_{rev} E(J)$ equations above, we obtain $\Delta_{std} E(J)$ = 0.89 and $\Delta_{rev} E(J)$ = 0.87. Thus, for the action-independent hypothesis:

$$\Delta_{std} X_i = 0.89 * w_{J,i} + w_{A_L,i} + w_{A,i} + w_{UI,i} \tag{29}$$
$$\Delta_{rev} X_i = 0.87 * w_{J,i} - w_{A_R,i} - w_{A,i} - w_{UI,i} \tag{30}$$

The results of a simulation based on this equation are plotted in the bottom of the second column from the left in Supplementary Fig. 1c, where it is still clearly possible to neurally separate judgment correlates despite their lower observed magnitude.

**(2)** High guess + lapse. Here we set $\gamma_{std} = \lambda_{std} = \gamma_{rev} = \lambda_{rev} = 0.5$, i.e., random licking in both blocks. Then according to the above equations $\Delta_{std} E(J) = \Delta_{rev} E(J) = 0$ and judgment correlates become impossible to observe:

$$\Delta_{std} X_i = w_{A_L,i} + w_{A,i} + w_{UI,i} \tag{31}$$
$$\Delta_{rev} X_i = -w_{A_R,i} - w_{A,i} - w_{UI,i} \tag{32}$$

**(3)** No lick/no-lick reversal. Here the mouse behaves the same way in reversed blocks as in standard blocks, which would mean that $\alpha_R \gg 0$, $\beta_R \ll 0$, and $|\alpha_R| \gg |\beta_R|$ (i.e., all stimulus present judgments are translated into right licks, and stimulus absent judgments into no licks). According to the above equations, this would mean that $\gamma_{rev} = \lambda_{rev} = 1$, and so $\Delta_{rev} E(J) = -1$. Meanwhile, as we changed nothing about the standard blocks, $\Delta_{std} E(J)$ is still 1. Thus:

$$\Delta_{std} X_i = w_{J,i} + w_{A_L,i} + w_{A,i} + w_{UI,i} \tag{33}$$

$$\Delta_{rev} X_i = -w_{J,i} - w_{A_R,i} - w_{A,i} - w_{UI,i} \quad (34)$$

As a result, judgment correlates, if they exist, are no longer positively correlated across the contrasts, and become fully entangled with the action-related correlates.

**(4)** Always air-licking on no-lick trials. Here we keep all parameters the same as in the negligible guess + lapse scenario, but replace no-lick trials with right licks in the standard block and left licks in the reversed block. Then the action terms cancel out, and we get both direction terms in each contrast:

$$\Delta_{std} X_i = w_{J,i} + w_{A_L,i} - w_{A_R,i} + w_{UI,i} * \Delta_{std} E[UI] \quad (35)$$
$$\Delta_{rev} X_i = w_{J,i} + w_{A_L,i} - w_{A_R,i} + w_{UI,i} * \Delta_{rev} E[UI] \quad (36)$$

Thus, judgment correlates, if they exist, become fully entangled with direction-related correlates. Note that for the purpose of the simulation in the right-most column of Supplementary Fig. 1c we also assume the magnitude of uninstructed movement differences across lick directions is small (as apparent from Fig. 1c) and can be neglected, but the judgment-direction entanglement will occur regardless.

In sum, even assuming mice solve the blocks task via learning a fixed rule based on lick port presence/absence, it is still possible to neurally separate action and judgment correlates. This separation relies on several assumptions which are empirically borne out by the data: (1) uninstructed movements are correlated with licking, (2) low guess and lapse rates, (3) successful reversal of lick vs. no lick responses to the same stimulus across blocks, and (4) negligible air licks. That said, the contrasts demonstration used here (Supplementary Fig. 1c) is idealized with pure selectivities — for mixed selectivities, this approach is less likely to work. Hence, we also employ linear regression (Figs. 3j, 4i) and population classification (Fig. 4j) in our neural analyses.
